# Genomic signatures of increasing disease burden in recent prehistory

**DOI:** 10.1101/2025.07.22.666087

**Authors:** Keith D. Harris, Yuval Talmor, Merav B. Yefe Nof, Nimrod Marom, Yitzchak Jaffe, Viviane Slon, Gili Greenbaum

**Affiliations:** Department of Ecology, Evolution and Behavior, The Hebrew University of Jerusalem, Jerusalem, Israel; Alpha Program, Future Scientist Center, The Hebrew University Youth Division, Jerusalem, Israel; School of Archaeology and Maritime Cultures, University of Haifa, Israel; Department of Anatomy and Anthropology and Department of Human Molecular Genetics and Biochemistry, Gray Faculty of Medical and Health Sciences, Tel-Aviv University, Israel; The Dan David Center for Human Evolution and Biohistory Research, Tel-Aviv University, Israel

## Abstract

One of the strongest selection pressures experienced by human populations is that driven by diseases on immune-related genomic regions. While it has been hypothesized for some time that disease burdens increased with the shift to agricultural and urbanized lifestyles, direct evidence for this hypothesis is lacking. Here, we capitalize on the accumulation of ancient genomic data to study changes in disease burden in human populations over the past 12,000 years. We investigated changes in genetic diversity and balancing selection in two distinct geographical and cultural centers in Southwest Eurasia and Eastern Asia, and found that not only is the major histocompatibility complex (MHC) the genomic region with the most substantial increases in diversity in terms of enrichment of genes and rates of increase, but that the rates of changes for individual genes are highly correlated between the two cultural centers, indicating similar selection pressures. We identify periods of time with substantial peaks in MHC diversity increase that primarily correspond to periods with settlement intensification, increased connectivity, and the expansion of animal domestication, which suggest that the most intensive disease burden occurred following the transition to sedentary lifestyle but prior to urbanization. These findings demonstrate the potential of our approach in uncovering the interplay between cultural shifts and selection, and provide strong support to the hypothesis that the levels of disease burden have substantially increased in recent prehistory following changes in lifestyle, connectivity and the introduction of domesticated animals.

## 1 Introduction

The recent surge in the availability of ancient genomes has revolutionized the way we study the past [1, 2]. This has been particularly true for the study of population movements and admixtures in recent prehistory [3– 5]; however, the study of selection processes using ancient DNA (aDNA) has proven challenging. While some cases of directional selection have been identified (e.g., the increase in lactase persistence alleles over time, presumably in relation to increased milk consumption; [6, 7]), they remain rare. The scarcity in identification of such cases may not necessarily be due to their absence, but rather a consequence of methodological biases and the limited availability of suitable ancient genomic data.

Human populations have experienced major transitions in lifestyle and living environments in recent prehistory, such as the transition to a sedentary lifestyle in the Neolithic and the later emergence of urbanized societies; these transitions generated selection pressures. One of the strongest types of selection pressures experienced by human populations is that driven by diseases on immune-related genomic regions [8], with specific diseases mainly inducing directional selection for particular resistance alleles, and overall disease burden inducing balancing selection that increases genetic diversity over larger genomic segments [9]. Although it has been hypothesized for some time that disease burdens increased as cultural complexity of human societies increased [10–12], direct evidence for this hypothesis is lacking.

Increases in disease burden, the cumulative fitness effect of pathogens on populations, could have been generated in several ways as new cultural forms emerged. For example, the growing proximity to domesticated animals likely led to increase in zoonotic spillovers [10, 13]. In addition, the transition to sedentary, agricultural, or urbanized societies was associated with increased population densities and more crowded living conditions [14], which would have likely increased the transmission rates of diseases. This would have allowed diseases that had previously been below the epidemic threshold (*R*_0_ = 1; [15]) to establish and become endemic. Some paleopathological findings [11, 16–18] are indeed suggestive of increased disease burden in the transition to sedentary agricultural societies, while other studies could not find support to the hypothesis [19–21]. However, inferring disease burden from skeletal remains is difficult because it is often unclear how to tease apart the health burden generated by lifestyle changes (e.g., diet, workload stress) from the burden inflicted by disease [22], and a high frequency of pathological findings may actually indicate low disease burden (the “osteological paradox”; [23]). Recent studies of ancient pathogen genomes have traced the origins and spread of several major diseases to the Neolithic period [24–29], but they cannot directly assess the burden these pathogens imposed on past populations.

Because the fitness cost generated by disease burden is tightly linked to immune-response, disease burden constitutes a strong factor in shaping immune-related genomic regions, particularly the major histocompatibility complex (MHC). The directional selection signature of some specific diseases can be observed in contemporary populations [30, 31]. However, genetic variation in immune-related genomic regions is mainly generated and maintained by balancing selection, because of (i) heterozygote advantage, where individuals with a larger immune-response repertoire can generate immunological responses to a wider diversity of pathogens, and (ii) frequency-dependent selection, where rare alleles are advantageous because pathogens are less likely to have evolved to evade immune responses involving them [9, 32]. Due to this balancing selection and the substantial (negative) contribution of disease environments to human fitness, the MHC is the most diverse region in the human genome [33]. This diversity is a response to the level of balancing selection induced by disease burden. Therefore, changes in disease burden levels may lead to changes in MHC diversity.

A promising direction for revealing selection processes is by tracking genetic changes over time using aDNA [34]. However, there are many challenges to this approach; for example, demographic processes and gene flow can obscure changes in allele frequencies, while sampling biases can affect the resolution and reliability of detected patterns. In addition, selection pressures are most often polygenic [30], and tracking changes in particular SNPs is often insufficient to reveal multi-locus signatures. Several studies have found evidence from aDNA of directional selection on variants related to diseases [34–41]. However, even in epidemics that presumably generated strong selection pressures, such as the Black Death in Europe, it is difficult to attribute selection to changes in variant frequencies [42]. Another challenge in studying diseases using aDNA is that many genomes are sequenced using a SNP panel that is biased, by design, towards high-frequency variants [43]. This panel is suited to demographic and population structure inferences, but less so to studying selection processes. Imputation tools can reduce this bias by expanding the set of variants beyond the SNP panel, and have shown some promise in enabling the study of multi-locus selection pressures in polygenic phenotypes [44, 45]. Although it is unclear whether imputed genotypes can be used to reliably estimate genetic diversity of ancient genomes, they can potentially enable tracking of changes in diversity and balancing selection in immune-related genomic segments.

Here, we study disease burden in human populations over time by tracking historical shifts in genetic diversity and balancing selection. We develop methods that can reliably estimate individual-level heterozygosity from ancient genomes, and implement them in genomic scans for balancing selection. Using these new methods, we study the last 12,000 years in two of the main geographical regions in which culture underwent major transitions interdependently, Southwest Eurasia and Eastern Asia, and identify periods with intensive disease burden. The new perspective on historical disease burden presented here can open new avenues for studying the genomic and cultural impacts of diseases on human evolution.

## 2 Results

### 2.1 Deriving unbiased heterozygosity estimates in aDNA using imputation

To identify selection by tracking changes in genetic diversity, heterozygosity must be reliably estimated in aDNA genomes. This is challenging owing to biases of commonly used SNP capture panels (e.g., 1240K), which are enriched for highly polymorphic variants [43]. Differences in sequencing strategies (i.e., shotgun sequencing versus SNP capture) further exacerbate the problem, by leading to substantial under- or overestimation of heterozygosity, masking the true levels of genetic diversity (Fig. S1). Finally, the low depth (<1x) of most aDNA sequencing libraries can limit heterozygosity calling, further reducing the ability to estimate the genetic diversity of genomic regions. A widely used approach for dealing with missing genotypes in genetic data is to impute genotypes using modern reference panels. While imputed genotypes generated from aDNA data can be useful for analyses of common variants [46], it is unclear whether the same imputed genotypes can be used to produce accurate estimates of heterozygosity for aDNA, and to what extent different sequencing strategies and read depths affect the bias in these estimates.

To generate accurate heterozygosity estimates from imputed genotypes in aDNA, we developed a pipeline called `impHet` that uses the imputed genotypes from the imputation tool GLIMPSE2 [47] to estimate heterozygosity for genomic windows (Fig. 1A). Because the imputation accuracy of GLIMPSE2 is limited to high frequency alleles (MAF *>* 0.01) in aDNA genomes [46], variants with lower MAF values are commonly filtered out. However, MAF-based variant filtering introduces a bias for higher heterozygosity, making it unsuitable for generating accurate estimates (Fig. S1). To address this, we instead filtered out genomes with low expected accuracy of their estimated heterozygosity. To determine expected accuracy, we designed a downsampling scheme that recapitulates biases in aDNA sequencing. First, we generated downsampled versions of high-coverage aDNA genomes so that their read distributions match those of low-coverage genomes from different sequencing strategies (Fig. 1B). This procedure of mimicking the read distribution of real aDNA sequences ensures that most biases generated by SNP panels and other limiting factors of aDNA sequencing are represented in the simulated downsampling. We used randomly-selected low-coverage genomes listed in the largest currently available aDNA dataset, the Allen Ancient DNA Resource (AADR; [2]), and generated 1000 downsampled versions for each high-coverage shotgun-sequenced genome (`SI table1.xlsx`). We then compared the heterozygosity estimates of the full high-coverage genomes with those of the simulated, downsampled versions of the same genome. The 1000 genomes used for mimicking read distribution in the downsampling procedure were chosen to be roughly representative of sequencing strategies and depths in the AADR, and include 50% genomes generated by SNP capture and 50% genomes generated by shotgun sequencing. We evaluated several high-coverage genomes with different levels of uracil-DNA glycosylase (UDG) treatments to examine its effect on the estimates.

**Figure 1:**
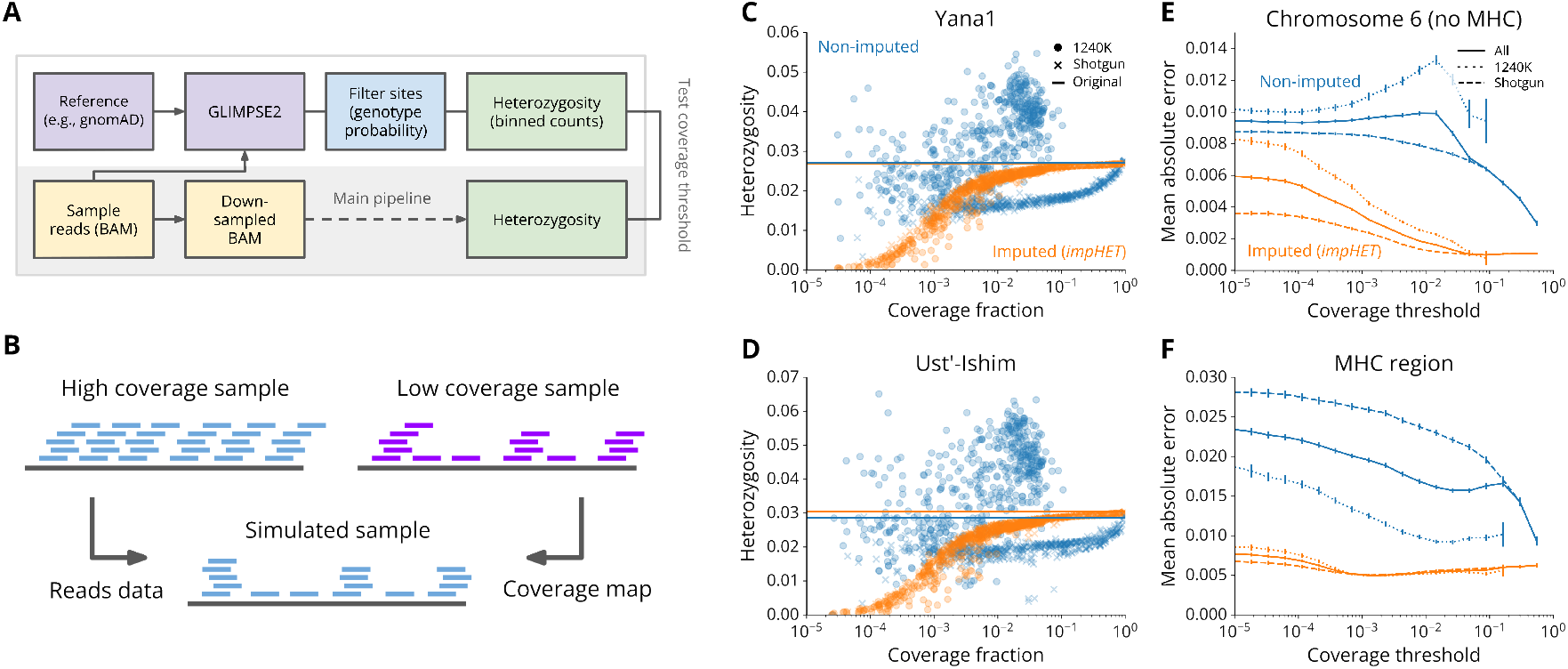
Evaluation of imputation-based estimation of heterozygosity in low-coverage aDNA genomes using impHet. (A) Pipeline for imputing heterozygosity, `impHet`. BAM files are processed by GLIMPSE2 using a phased reference panel (gnomAD v3.1.2) to generate imputed genotypes, which are then filtered and summed in binned genomic regions (either fixed genomic windows or gene boundaries). (B) To evaluate the estimate accuracy, a BAM file of a high-coverage genome is downsampled using this scheme, and processed by `impHet` to compare to the results of the full coverage BAM. This reproduces the effects of biases in different aDNA sequencing methods by recapitulating low-coverage sequencing read distributions. Using a high-coverage genome ‘source’ and low-coverage genome as a ‘template’, we resample reads from the source genome to generate realistic read distribution panels using multiple templates. These replicates can then be compared to the “ground truth” estimate of heterozygosity (the heterozygosity of the source genome). (C–D) Heterozygosity estimates either using `impHet` (orange) or from non-imputed (blue) genomes generated by downsampling the Yana1 (C) and Ust’-Ishim (D) genomes. 1000 low-coverage read-distribution templates were used for each panel. Coverage fraction relates to the proportion of variants with a read depth ≥ 4 in each downsampled genome, out of all variants in the panel for the specific genomic region. The lowcoverage genomes used as templates were selected randomly from the AADR to reflect commonly available sequencing strategies and depths. Horizontal lines indicate the “ground truth” heterozygosity for the full coverage of the source genome, either non-imputed (blue) or from `impHet` (orange). (E–F) Comparison of estimates of heterozygosity for different coverage thresholds using `impHet` (orange) or non-imputed (blue) estimates, in chromosome 6 (E) and the MHC (F). Eight high-coverage source genomes and the same 1000 low-coverage read-distribution templates used in (C–D) were used in these analyses. Coverage relates to the proportion of variants with a read depth ≥4 in each downsampled genome, out of all variants in the panel for the specific genomic region. The mean absolute error is the average distance in heterozygosity from the ground truth heterozygosity, for all downsampled genomes that have coverage above the specified threshold. Error bars indicate the 95% confidence intervals for each threshold, generated by bootstrapping. The value is shown separately for SNP capture sequencing (dotted), shotgun sequencing (dashed) and both sequencing methods (solid line).

We found that non-imputed heterozygosity is inaccurate, is inconsistent across different coverages, and produces different values for different sequencing strategies compared to the heterozygosity of the full highcoverage genomes (blue circles and crosses in Fig. 1C–D and Fig. S2; see also Fig. S1). SNP capture read distributions (e.g., the 1240K SNP panel) lead to over-estimated heterozygosity (circles in Fig. 1C–D and Figure S2), while shotgun read distributions lead to under-estimated heterozygosity (crosses in Fig. 1C–D and Fig. S2). On the other hand, heterozygosity estimates generated by `impHet` have a higher accuracy (orange circles and crosses in Fig. 1C–D and Fig. S2). Notably, the estimates for both SNP capture and shotgun sequencing converge to the full coverage value with increasing genomic coverage (solid blue in Fig. 1C–D and Fig. S2. Our imputation-based estimate has substantially less error compared to the nonimputed heterozygosity estimates (Fig. 1E). In the MHC, which has higher recombination and is expected to have lower accuracy for imputed genotypes [9], our estimate is also more accurate than the non-imputed heterozygosity, and consistent between sequencing strategies (Fig. 1F; Fig. S3). We observe that for coverage *>* 2% the error in the `impHet` estimate of heterozygosity is below 0.002 for chromosome 6, and below 0.01 for the MHC (orange lines in Figure 1E–F); the ~ 5 times higher error in the MHC may be attributable to the ~ 3–5 times higher heterozygosity levels in the MHC (Figs. S2 and S3). In addition, our estimate has lower variance compared to the non-imputed heterozygosity (indicated by the distribution of downsampled genomes in Fig. 1C–D). This demonstrates that, when filtering for genomes with coverage *>* 2% `impHet` can derive reliable estimates of heterozygosity.

### 2.2 The MHC is a focal genomic region for increases in genetic diversity

In our analyses, we mainly consider two sets of ancient genomes from the AADR dataset [2], each defined by having a shared ancestry component and geographic continuity, broadly representing two distinct cultural centers (Fig.S4): a Western group in Western Asia and Southeastern Europe (523 individuals), and a Eastern group in Eastern Asia (178 individuals) spanning the last 12,000 years. To identify genomic regions that have experienced changes in genetic diversity, we developed a simple statistical approach. For a given set of genomes, we first estimated the heterozygosity of each genome in the genomic region of interest using `impHet`. Applying a simple linear regression, we then tested whether any significant change in heterozygosity over time occurred. We consider the slope of the regression, denoted as *β*_*H*_, as the rate of change of heterozygosity in this genomic region. For example, we detected a significant (*p* < 10^−^13) change of heterozygosity in the MHC in the Western group over the last 12,000 years, with a positive rate of *β*_*H*_ = 0.00278 heterozygosity increase per 1000 years (Fig. 2A). Strikingly, this rate is 16 times higher than the genome-wide rate of increase in heterozygosity over this time period for this group (Fig. S5A–B).

**Figure 2:**
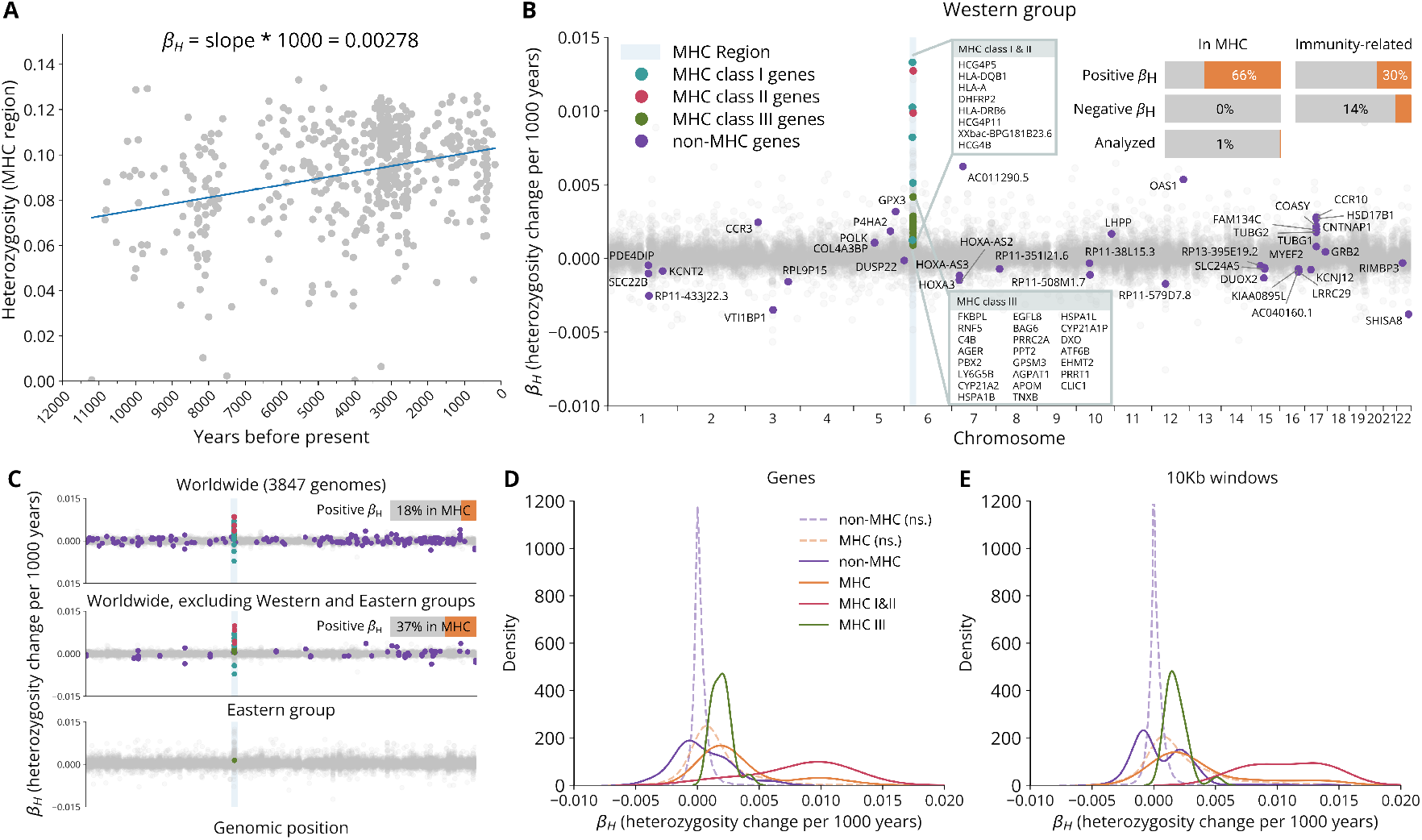
Genome-wide scan for changes in heterozygosity over time. (A) The computation of *β*_*H*_ for MHC heterozygosity in the Western group. Gray dots show MHC heterozygosity estimates for individual genomes in the Western group. The blue line indicates the slope of the linear regression. At the top of the panel, the computation of *β*_*H*_ is illustrated for this regression, yielding a value of an increase in heterozygosity per 1000 years of 0.00278. (B) Results of the genomic scan in the Western group using *β*_*H*_ for genes, ordered by genomic position. The MHC region is highlighted in blue. Significant genes are colored turquoise for MHC class I, magenta for MHC class II, green for MHC class III and purple for all other genes. Non-significant genes are colored gray. All significant genes are annotated, with boxes for the densely clustered MHC genes. In the top right corner are bars showing enrichment of MHC and immune-related genes in the significant hits of the genomic scan. The results are separated to positive (increase in heterozygosity) and negative (decrease in heterozygosity) trends. The bottom bar shows the proportion of genes in the MHC out of all those genes that we analyzed, for comparison. (C) Genomic scans for all genomes in the AADR that passed the coverage threshold (top panel), all genomes that passed this threshold excluding the Western and Eastern groups (middle panel), and for the Eastern group (bottom panel). The MHC region is highlighted in blue, and significant genes are colored according to their genomic position as in (B). The proportion of significant genes with positive *β*_*H*_ values in the MHC is indicated in the top right of each panel. (D) Kernel density estimates for *β*_*H*_ values for significant MHC genes (orange), MHC class I&II genes (magenta), MHC class III genes (green) and non-MHC genes (purple), values of which are shown in (B). Kernel density estimates for non-significant genes in MHC and non-MHC are represented by orange and purple dashed lines, respectively. (E) Kernel density estimates for *β*_*H*_ values from a genomic scan using fixed-size windows of 10Kb. Shown are significant MHC genes (orange), MHC class I&II genes (magenta), MHC class III genes (green) and non-MHC genes (purple). Kernel density estimates for non-significant genes in MHC and non-MHC are represented by orange and purple dashed lines, respectively.

We then apply this approach as a genomic scan, computing *β*_*H*_ across all genes, or across 10Kb genomic windows. In these genomic scans, we consider the strength of the signal in terms of the magnitude of change detected (the values of *β*_*H*_); after correcting for multiple testing, we retain only those genomic regions that achieve significance in at least 95% bootstrap replicates. In the Western group, where we have more genomes and therefore stronger statistical power (Fig. S4), we found 71 genes that showed a significant change in heterozygosity (Fig. 2B), 66% of them with a positive change and 34% with a negative change (for the full list of genes, see Tables S1 and S2). While some of these hits are clustered along the genome, we found relatively low linkage disequilibrium (LD) among significant genes in the MHC (Fig. S6). Surprisingly, out of those with a positive change, more than 60% were found in the MHC (Fig. 2B), even though MHC genes constitute less than 1% of the genes we tested. Genes with immune-related function are enriched for significantly positive *β*_*H*_ values, but to a lesser extent than the MHC itself (Fig. 2B). All genes with significant change in the MHC had positive *β*_*H*_ values (Fig. 2B), and these values were substantially higher in the MHC compared to the rest of the genes (two-sided Wilcoxon rank-sum test, *p*-value < 1 × 10^−4^; 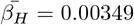 for MHC genes compared to 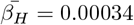 for non-MHC genes; Fig. 2D). This trend was also observed in (i) a scan using all 3847 individuals in the AADR that passed our coverage threshold (top panel in Fig. 2C), (ii) a scan using these individuals but excluding the Western and Eastern groups (middle panel in Fig. 2C), (iii) a scan using the Eastern group (bottom panel in Fig. 2C; Fig. S8B), (iv) a scan using 10Kb genomic windows (Fig. 2E), and (v) a scan using a different gene annotation release (Fig. S9).

The MHC is typically divided into three genomic regions (classes), with MHC class I broadly related to viral responses, MHC class II to bacterial and fungal responses, and MHC class III associated with inflammatory and less-specific immune functions [48] (some functional crossover occurs between classes). The MHC genes with the largest *β*_*H*_ values are found in MHC classes I and II and, although more numerous, significantly increasing genes in MHC class III had lower rates of increase (Fig. 2B,D). For genes outside the MHC, some of those increasing in heterozygosity were also related to the immune system (e.g., OAS1, CCR3, CCR10), while others are related to other systems, such as DNA polymerization (e.g., POLK) or oxidative stress response (e.g., GPX3) (Table S1). While there were no genes that decreased in heterozygosity in the MHC, 24 genes outside the MHC did show a significant reduction in heterozygosity, possibly due to directional selection (Fig. 2B–D; Table S2).

In the Eastern group, we also detected an overall significant increase in heterozygosity in the MHC, with a similar magnitude of *β*_*H*_ = 0.00275, which is 7 times higher than the whole-genome rate of increase for this group (Fig. S5C–D). Unlike in the Western group, we detected only one gene with significant changes in heterozygosity, EGFL8 in MHC class III (Fig. S8), most likely due to the smaller sample size of the Eastern group (Fig. S4). Nevertheless, the EGFL8 gene is also significant in the Western group genomic scan, and, when conducting the analysis on non-significant genes, in the Eastern group the *β*_*H*_ values in the MHC were higher compared to other genomic regions (Fig. S8E).

Overall, our results show that in both geographic regions the increase in genetic diversity in the MHC is substantially higher than the rest of the genome (Fig. S5). This suggests that the MHC is the genomic region that has experienced the largest increase in genetic diversity over the last 12,000 years, likely driven by immune-related balancing selection.

### 2.3 Substantial increase in MHC class II heterozygosity around 7000BP

To further characterize the process of increase in MHC diversity, we tracked the average heterozygosity in overlapping 2000-year sliding windows over this period in the two groups (Fig. 3A–B and Fig. S10A–B). To determine whether substantial changes in diversity occurred in specific time periods, we performed regression analyses in overlapping 3000-year sliding windows (we used a larger sliding window for regression analyses to capture periods of change rather than a snapshot of heterozygosity), and computed the *β*_*H*_ statistic for each window separately (Figs. 3C–D and S10C–D). Finally, to contextualize the increase in heterozygosity in terms of increase in balancing selection, we used a simplistic mathematical model to derive balancing selection coefficients for each 3000-year window (Figs. 3E–F and S10E). These coefficients should be interpreted as a simple quantification of the additional level of balancing selection experienced by the population in a given time interval over background levels of balancing selection, mutation and drift (see *Methods*). To relate this signal to other genomic regions, we compared the MHC class signals to the distribution of balancing selection coefficients in all genomic windows of a similar scale (1Mb) outside the MHC (Figs. 3E–F and S10E).

**Figure 3:**
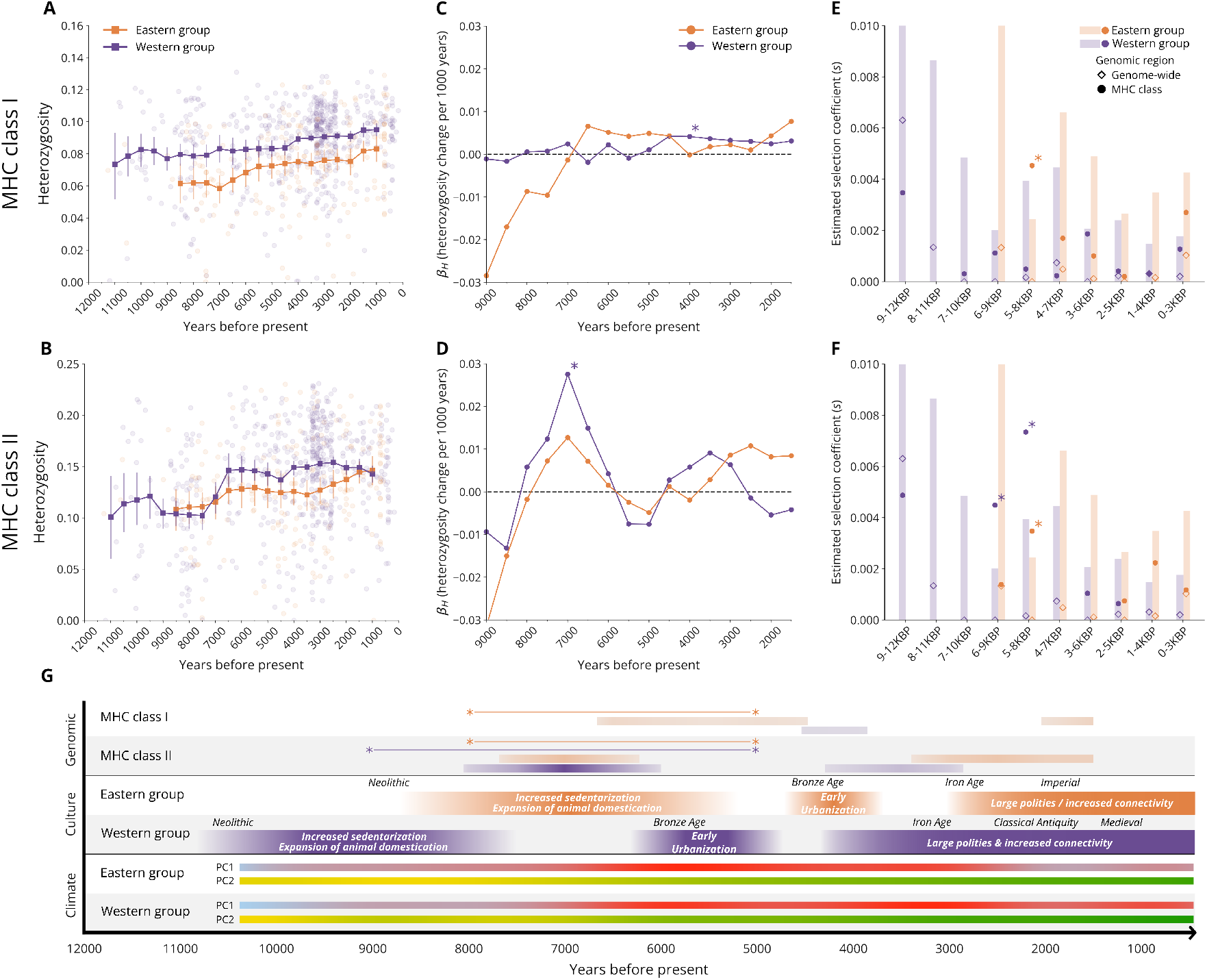
Temporal signals of balancing selection in MHC class I and MHC class II. (A–B) Sliding window analysis of heterozygosity of MHC class I (A) and MHC class II (B), in the Western (purple) and Eastern (orange) groups. Solid lines with squares represent mean heterozygosity calculated over 2000-year windows centered around each window midpoint. Circles represent heterozygosity values of individual genomes. Error bars indicate the 95% confidence intervals of the mean, generated using bootstrapping. Windows with less than 10 genomes are not shown. (C–D) Temporal analysis of *β*_*H*_ in MHC class I (C) and MHC class II (D) in the two groups. Solid lines with circles represent *β*_*H*_ values calculated over 3000-year windows centered around each window midpoint. Asterisks indicate significant regressions for the specific window. (E–F) Estimates of balancing selection in MHC class I (E) and MHC class II (F) in the two groups. Each column represents an overlapping 3000-year period. Shaded bars represent the extent of the upper 95% confidence interval of the genome-wide distribution of *β*_*H*_ values. Empty diamonds represent the mean genome-wide signal, and filled circles represent the signal of the specific MHC class. Windows in which the MHC class signal is outside the confidence interval of the genome-wide signal are indicated with asterisks. (G) Cultural and climate context of the genomic signals we identified. The top section shows the main genetic signals from panels C–F: horizontal gradient bars are visualizations of the signal from panels C–D (darker colors represent higher *β*_*H*_ values); asterisks connected by lines are visualizations of the temporal windows where a significant signal of balancing selection was identified in panels E–F. The middle section shows a timeline of archaeological periods (labels at the top) and cultural processes that could be related to changes in disease burden (gradients with overlaid text). The bottom section shows the two main PCs of a set of climate variables in the two geographic regions. PC1 mainly captures temperature variables, from blue (colder) to red (hotter), and PC2 mainly captures precipitation variables, from yellow (drier) to green (wetter).

We observe that, although MHC diversity increased in both groups and in all three MHC classes (Figs. 3A– B and S10A), shorter periods of dramatic increases can be observed. Most strikingly, the period around 7000BP shows a significant increase in MHC class II genetic diversity, in both groups (Fig. 3B). In both geographic regions, this period is characterized by increased sedentarization and expansion of animal domestication, and is earlier than the emergence of urban centers (Fig. 3G). The substantial increase in MHC class II diversity during this time is also demonstrated by a peak with the highest *β*_*H*_ values, both in the Western and Eastern groups, in the temporal window centered on 7000BP (Fig. 3D). The changes in genetic diversity in MHC class II appear coordinated around this peak, with positive and increasing rates from about 7500BP to 7000BP, and decreasing rates from 7000BP to about 4500BP. In the Western group, the increase in diversity is particularly rapid, with significant *β*_*H*_ rates of 0.0275 (*p*-value < 1× 10^−5^). This pattern is also reflected in the balancing selection coefficients, where we observe high coefficients (*s* = 0.004 and *s* = 0.007 in the Western group and *s* = 0.003 in the Eastern group) in the windows centered around 7500–6500BP, which are significantly higher than the genome-wide distribution of coefficient values (Fig. 3F). Around this time point, 7000BP, we also observe relatively high peaks in rates of diversity increase (Fig. 3C–D) in the Eastern group in MHC class I, and in the Western group in MHC class III, but at lower rates (*β*_*H*_ =0.0066 in the Eastern group MHC class I at 6500BP and *β*_*H*_ =0.0046 in the Western group MHC class III at 7000BP). In the Eastern group MHC class I, the balancing selection coefficient around 6500BP is significantly above the genome-wide distribution (*s* = 0.005; Fig. 3E). Notably, these increases in MHC diversity cannot be explained by demographic processes, because no substantial increase in genome-wide diversity is observed in this time period in either group (Figs. S10D and S10D).

For all MHC classes, we generally observe higher levels of heterozygosity in the Western group than in the Eastern group, where the gap is maintained throughout the process (Figs. 3A–B and S10A). This gap in diversity disappears in the MHC classes II and III as we approach modern times, whereas in MHC class I a gap remains (this gap is observed also in modern populations; Fig. S11). In MHC class I, we observe a relatively constant and steady increase in diversity, that perhaps started later in the Eastern group than in the Western group, with increases in diversity starting only around 8500BP in the Eastern group (Fig. 3C). In MHC class II, however, we observe two peaks of increase in diversity: the first is the coordinated peak around 7000BP, and the second, less substantial than the first, occurring 4000–3000BP in the Western group, and later over the last 3000 years in the Eastern group (Fig. 3D). In both groups, the peaks were separated by a period of constant levels of heterozygosity, and even slight decreases around 5500-5000BP. In MHC class III, we observe a period of increase in diversity in the Western group at around 8000–7000BP, and in the Eastern group the increase occurred mainly over the last 3000 years (Fig. S10C), similarly to the later peak observed in MHC class II in this group (Fig. 3D). Overall, our detailed characterization of the temporal dynamics of diversity in MHC classes enables a nuanced understanding of how immune-related selection pressures unfolded over time. This reveals a substantial, shared peak in disease burden following the transition to sedentary life-styles and animal domestication, and prior to urbanization, as well as region-specific patterns that correspond to cultural transitions across Southwest Eurasia and Eastern Asia.

### 2.4 Selection on genes in the MHC is highly correlated between the Western and Eastern groups

Our analyses indicate that immune-related genetic diversity increased worldwide, and specifically in two distinct geographic regions, that of the Western group and that of the Eastern group. These changes may reflect increase in disease burden in the two regions following major cultural transitions, but it is unclear whether the processes in the different regions generated selection pressures on the same genes in the two groups, or rather on different sets of genes. To examine this question, we evaluated the correlation between the rates of change in heterozygosity for particular genes (i.e., *β*_*H*_ values), in the two groups (Fig. 4A). A high correlation would indicate that genes that increase in diversity in one group also tend to increase in diversity in the other. We evaluated this correlation for MHC genes and non-MHC genes, and we separated the analysis to genes for which significant *β*_*H*_ values were attained (in either group) and those without significant changes, for an additional control.

**Figure 4:**
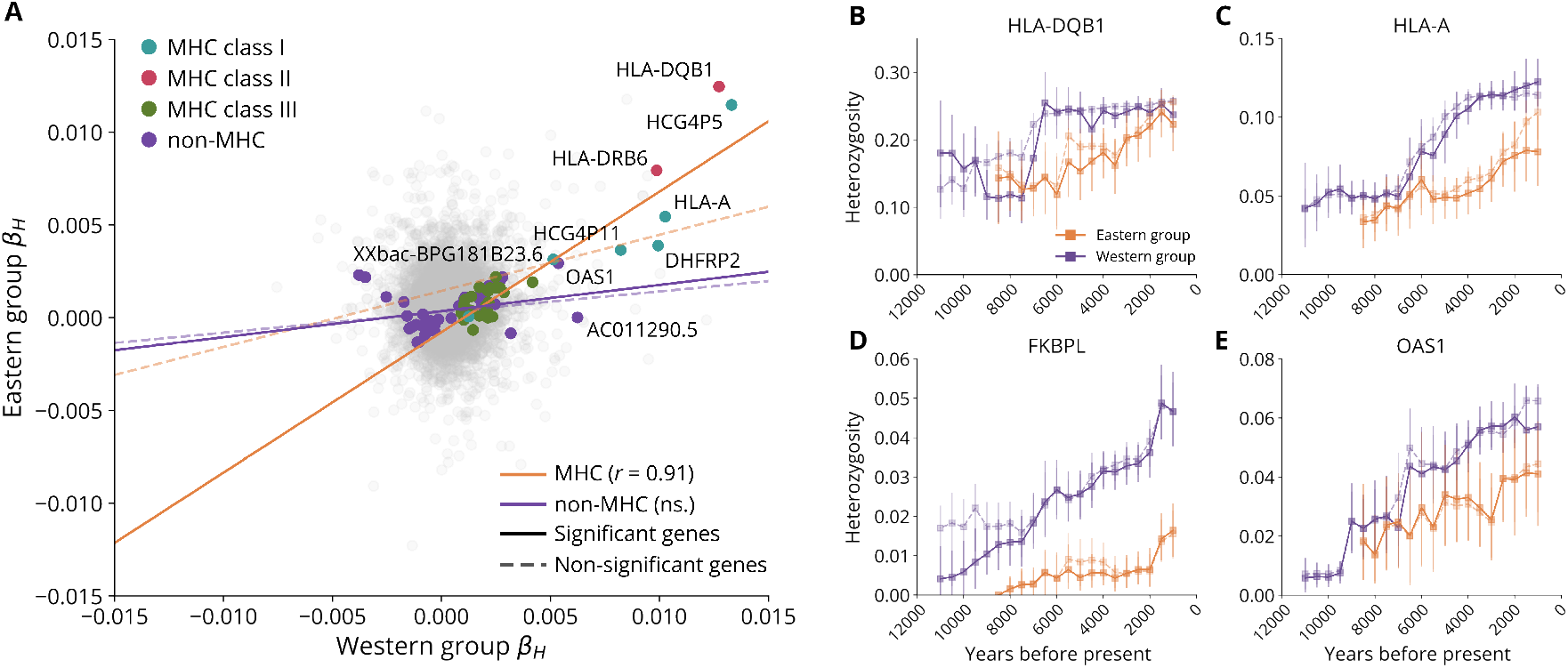
Correlation of heterozygosity changes between the Western and Eastern groups. (A) Correlation of *β*_*H*_ between groups. Each dot represents a gene, with the value of *β*_*H*_ for the Western group on the x-axis, and the value for the Eastern group on the y-axis. Genes with significant results in either group are colored according to the genomic region: turquoise for MHC class I, magenta for MHC class II, green for MHC class III and purple for all other genes. Genes with no significant result are colored gray. Genes with | *β*_*H*_ |*>* 0.005 are annotated. The solid orange line is the regression result for MHC genes with a significant regression, and the dashed orange line is the result for MHC genes with no significant regressions. The solid purple line is the result for genes outside the MHC with a significant regression, and the dashed purple line is the result for genes outside the MHC without a significant regression. (B–E) Sliding window analysis of temporal trends of heterozygosity in the Western and Eastern groups for HLA-A (B), HLA-DQB1 (C), FKBPL (D) and OAS1 (E), significant genes that were identified by our *β*_*H*_ genomic scan. In addition to average observed heterozygosity (solid lines), we analyzed expected heterozygosity of each window (dashed lines).

We observe a strong correlation (Pearson coefficient *r* = 0.91, *p*-value < 1 × 10^−10^) between rates of significant increase in diversity in the MHC genes when comparing the Western and Eastern groups (Fig. 4A). In contrast, we find no significant correlation for non-MHC genes that have significant *β*_*H*_ values. Genome-wide increases in diversity, as well as gene flow between the groups, would be expected to generate a statistical correlation for comparison of genes; in this regard, the correlation of non-significant non-MHC genes can be treated as a control that, at least partially, captures effects unrelated to selection. We observe a weak but significant correlation for non-significant non-MHC genes (*r* = 0.08, *p*-value < 1 × 10^−36^), with a smaller slope (0.11 compared to 0.76 for significant MHC genes), suggesting that the strong correlation in the MHC cannot be explained by these confounding factors alone, and therefore likely reflects similar selection pressures on MHC genes in the two groups. In the MHC, the signal of both significant and nonsignificant genes was correlated (Fig. 4A), but the correlation is substantially stronger for the genes where the increase was statistically significant compared to those where it was not (*r* = 0.91 compared to *r* = 0.24, respectively), indicating that the correlation between groups is specifically high for those genes we highlighted as experiencing balancing selection (Fig. 2B).

The strong correlations we find here indicate that particular genes were under balancing selection in both groups in this time period. For example, the gene for which we detected the largest increase in diversity in the Western group, HLA-DQB1, is also that with the highest increase in diversity in the Eastern group (*β*_*H*_ =0.0127 in the Western group and *β*_*H*_ =0.0125 in the Eastern group). Notably, specific variants in HLA-DQB1 were identified by a number of previous studies using tests for directional selection [41, 45, 49], which could be a reflection of negative-frequency-dependent balancing selection on the gene. When considering the temporal changes in diversity for this gene (Fig. 4B), we observe that they are similar to the overall trends for MHC class II (Fig. 3D), in which this gene is located, with a dramatic increase of diversity in the Western group at about 7000BP. For the top hit in MHC class I, HLA-A (HCG4P5, HLA-complex Group 4 Pseudogene 5, is nested within HLA-A), a well-studied gene related to viral immune defense, the temporal trends (Fig. 4C) are also similar to the broader patterns observed for its MHC class (Fig. 3C), where diversity is higher in Western group throughout the time period, and increases in diversity in the Eastern group occur later than in the Western group. The top hit in MHC class III, FKBPL, shows an increase over time in both groups, but with trends somewhat different than those of MHC class I and MHC class II (with a larger gap between the two groups that increases towards the present), highlighting that these differences may reflect different selective pressures for the different functions of the MHC classes (Fig. 4D).

Examining the temporal trends in genetic diversity of particular genes can provide insights into evolutionary processes. For example, when considering a non-MHC immune related gene that has been identified as significantly increasing in diversity in the Western group, OAS1, we observe patterns similar to those of HLA-A and MHC class I (Fig. 4E). OAS1 is a well-established antiviral factor, and also displays antibacterial activity [50]. In addition, the OAS locus shows adaptive introgression from Neanderthals [51]. Another non-MHC gene with an established role in immunity identified by our genomic scan is CCR3. CCR3 shows an increase in diversity around 7000BP in the Western group, similarly to MHC class II gene HLA-DQB1, and another increase from 4000BP, but a limited increase in the Eastern group (Fig. S13). CCR3 is a chemokine receptor that is linked to parasitic worm (helminth) infections [52], which likely became more common with the transition to sedentary lifestyles through the increased exposure to contaminated water and soils and to domesticated animals [53]. Additionally, selection pressures induced by helminths may have been longer-acting than those generated by other pathogens [31]. Therefore, it is possible that the signal we detected in this gene is related to the same underlying ecological and cultural processes as MHC genes (although there is no direct evidence that CCR3 polymorphism is advantageous). We also detected few strong heterozygosity-increase hits in non-immune genes, such as GPX3, which encodes glutathione peroxidase; however, here we do not observe a correlation between the groups, with an increase in diversity at earlier periods than in the MHC in the Western group (12000-7000BP), and a slight decrease in the MHC in the Eastern group (Fig. S13). These examples highlight the potential of tracking temporal levels of diversity at the levels of genes for revealing important details about changes in genetic diversity.

## 3 Discussion

We studied changes in disease burden in human populations over the past 12,000 years by examining changes in genetic diversity in immune-related genomic regions. We developed and validated a new approach to measure heterozygosity in ancient genomes, `impHet`, and used it to quantify changes in immune-related genetic diversity in two groups that represent distinct cultural centers. We performed genomic scans and quantified the increase in diversity across the genome over time, in the two focal groups and in worldwide groups including thousands of genomes, identifying the MHC as the genomic region with the most substantial increases, both in terms of enrichment of genes that increase in diversity and the rates of increase (Fig. 2). These findings provide a unique insight into changes in the levels of disease burden over recent prehistory, and their temporal coincidence with changes in lifestyle, connectivity and domestication of animals. The increase in diversity in specific genes is strikingly correlated between the two groups we examined (Pearson *r* = 0.91; Fig. 4), suggesting that the two cultural centers have experienced similar balancing selection pressures at the level of genes. Because our approach allows us to identify specific timings for signals of balancing selection at different genomic regions, the temporal signals we identified at immune-related genomic regions could be used to characterize the connection between major transitions and processes in human culture and associated changes in disease burden (Fig. 3G).

The observed changes in heterozygosity can be associated with large-scale cultural processes in both the Eastern and Western groups we studied (Fig. 3G). The rising rates of immune-related diversity from 8500BP in the Western group, peaking at around 7000BP, are loosely correlated with the spread of agriculture and domesticated animals from its Middle Eastern origins into Southeastern Europe, which entailed demographic expansion and contact between hunter-gatherers, farmers, and livestock [54, 55]. The later periods in which we observe increased rate of change in MHC diversity are at about 5000BP, following the emergence of large settlements during the 6^th^ millennium BP in a complex and nuanced “urban revolution” process [56, 57].

Both periods of increase in immune-related diversity, therefore, correspond to key expansion, aggregation and, importantly, connectivity events that can be expected to have resulted in increased exposure to pathogens, increased population densities, and increased disease transmission rates. A recent study that screened for ancient pathogens in aDNA libraries showed increased pathogen detection at around 5000BP in Europe [29]; this largely corresponds to the timing of disease burden increase we describe here, perhaps to the larger peak at 7000BP, with delayed arrival of diseases to Europe from the cultural centers of the Levant possibly explaining the later peak of pathogen observations in Europe. While ancient pathogen studies can provide much details on the emergence of diseases, they cannot quantify the burden diseases generate, and therefore ancient pathogen and our host-immunity approaches are especially complementary.

The genomes in the Eastern group are from the core of the cultural center of East Asia, as well as from its periphery (e.g. Mongolian Plateau, Eastern Kazakh Steppe, Tibetan Plateau, Korean Peninsula; Fig. S4). In this geographic region, we also observe a peak in MHC class II diversity change at around 7000BP, during a time when agriculture is becoming commonplace in late Neolithic villages across China [58], while in more northern regions, hunter-gathering communities persisted longer and eventually shifted towards pastoral economies [59]. From the 9^th^ millennium BP, increased sedentism and broader food exploitation spectrums have been documented in these areas [60]. As in the Western group, these cultural changes may have resulted in an increase in disease burden, reflected in significant balancing selection signatures in both MHC class I and MHC class II (5–8KBP in Fig. 3E–F). Later, in the 4^th^–3^rd^ millennium BP along the Yellow and Yangzi River valleys complex polities sprouted large urban complexes, supported by intensive agricultural economies, first of the Shang and Zhou dynasties and finally with the establishment of the first Qin empire. In the Eurasian steppe, this period is characterized by pastoralism with varying degrees of agricultural cultivation [61], at a time when the whole region saw intensified long-range interactions, trade networks and movement across vast landscapes [62, 63]. Therefore, the combination of increased connectivity in the northern part of the Eastern group with the rise of large polities in the southern part may have contributed to the signals of increased MHC diversity at the period of 3000-1500BP.

In addition to examining the association between immune-related diversity changes and cultural processes, our study adds to the rapidly growing set of loci that are candidates for selection in recent human evolution [34, 45, 49]. Some of the highest *β*_*H*_ hits from our genomic scan overlap with loci identified by previous studies using different tests for selection (summarized in Tables S1 and S2). For example, the HLA-DQB1 gene, which has the highest *β*_*H*_ value in both of the groups we studied, was also highlighted by previous studies [34, 45, 49]. This concordance between different selection tests is expected particularly in genes where many variants are under selection, because a balancing selection signal over a genomic segment is consistent with directional selection on multiple variants in this segment. The agreement between our results and other approaches for certain loci, along with the additional loci detected by our genomic scan, is particularly encouraging since it suggests that these different approaches can complement each other to identify strong candidate genes that have undergone selection during recent human evolution. Of the novel loci we identified, many have known immune functions or are in the MHC (Tables S1 and S2). Notably, MHC genes were strongly correlated between the two culturally distinct groups, while other genes showed different trajectories that could indicate different selection pressures in each group (Fig. 4).

While our approach enables tracking heterozygosity changes, identifying genomic regions of interest, and relating shifts in rates of change to historical periods, directly associating these signatures with cultural changes is difficult. In particular, given the sparsity of aDNA in periods and geographical regions relevant to studying major cultural transitions, it is not currently possible to evaluate how genomic signatures respond to various changes in disease burden, and it is expected that selection signals will lag after the appearance of their cultural and ecological drivers in the archaeological record. In addition, demographic processes and gene flow events can obscure selection signals; this is especially problematic considering that many periods of cultural change involved substantial migration and changes in population structure. This also means that care is required in evaluating how populations are defined; for example, we identified several genomes from the Iron Gates gorge region with differing ancestry profiles, the inclusion of which gives a strong signal for a single gene (ORD10AD1), but this signal was filtered out by our robustness test (Fig. S15). Due to these limitations, we can currently only provide a rough sketch of possible associations between disease-burden-enhancing cultural processes and immune-related genomic signatures. This is compounded by technical issues, such as the biases inherent in sequencing strategies that are widely used in generating aDNA libraries. The common SNP panels employed are enriched for highly polymorphic sites, rendering estimates derived directly from this data biased (Fig. S1). Our extensive validation of the `impHet` pipeline directly addresses these concerns, and supports the feasibility of using imputed genotypes to provide consistent estimates of heterozygosity for aDNA genomes with different sequencing strategies and depths.

Our findings here present compelling evidence that disease burden had indeed increased in the Neolithic in two major cultural centers, and that this rise was sufficiently substantial to trigger evolutionary processes that have shaped our genomes. We also suggest that this process was not temporally homogeneous, potentially peaking following major cultural transitions. Our study, therefore, highlights the potential of ancient genomics to address key questions in biological and cultural evolution, and to contextualize the archaeological record in terms of epidemiological and ecological processes.

## 4 Methods

### 4.1 Data curation

To track historical shifts in genetic diversity and balancing selection, we selected sets of ancient genomes with sufficient coverage for our heterozygosity estimating pipeline `impHet`, and that have a shared ancestry component and geographic continuity. To identify genomes matching these requirements, we analyzed the coverage of a large set of published ancient genomes from the AADR [2]; genomes passing our coverage threshold were then filtered based on their corresponding geographical position and ancestry. These genomes were then processed by `impHet` to generate heterozygosity estimates.

`impHet` is based on the GLIMPSE2 imputation pipeline [44]. GLIMPSE2 requires a phased reference panel for imputation, which determines the set of variants of the imputed genomes; the same reference panel is used to generate non-imputed genotypes when assessing the accuracy of imputation. We used the phased gnomAD v3.1.2 reference panel [64], which includes 4091 genomes from the 1000 Genomes Projects and Human Genome Diversity Project. As the panel is released for `hg38` and contains multiallelic variants incompatible with GLIMPSE2, we performed the following pre-processing steps before using the panel for any analyses. We first removed non-biallelic and non-SNP variants from the original panel. The filtered panel was lifted to `hg19` using the `bcftools +liftover` plugin [65]. The output was then normalized with `bcftools norm`, and again filtered for non-biallelic variants or any duplicate coordinates introduced during the lift-over process. The final panel included 66,024,110 variants from all autosomes.

To generate coverage estimates on our reference panel for genomes in the AADR, we first downloaded all publicly available BAM files associated with genomes published in the AADR v62 dataset [2]. We then assessed coverage on chromosome 6, and specifically in the MHC, against the normalized gnomAD reference panel using `bcftools mpileup`. We measured coverage as the fraction of variants in the normalized gnomAD reference panel that had a depth of at least 4x in the specific genome. Following the determination of the coverage threshold shown in Figure 1E–F and explained below, we filtered out genomes with less than 2.5% coverage on the gnomAD panel in the MHC region (this removes shotgun sequenced genomes that have good genome-wide coverage but low coverage in the MHC).

To extract sets of genomes from those that passed our coverage filter, we used the DORA interactive aDNA platform [66], which visualizes multiple layers of metadata associated with genomes in the AADR v62 dataset [2]. Using the ADMIXTURE model of ancient genomes in the AADR (*K* = 3) that is loaded into DORA by default, we selected geographical boundaries that had a shared ancestry component, and roughly overlapped two distinct geographical regions that correspond to two main cultural centers: a ‘Western group’ and ‘Eastern group’ (Fig. S4, `SI table1.xlsx`). Finally, we removed a small number of individuals from the Eastern group that were outliers in terms of their ancestry composition. In the Western group, we retained a subset of 29 individuals from the Iron Gates gorge region, which reflect in their distinct ancestry composition the complex interaction between genetic and cultural exchange at this crucial entry point to Europe before and during the Neolithic transition [67]. We also tested the effect of their exclusion from analyses on our results (Fig. S15).

After these filtering steps, the datasets for each group included 523 individuals in the Western group and 178 individuals in the Eastern group (individuals from each group are listed with metadata from the AADR in SI table1.xlsx). In cases where the AADR listed two separate genomes (from different studies) for the same individual, we used the genome with higher coverage.

### 4.2 Estimating heterozygosity with impHet

To answer the need for an unbiased estimator of heterozygosity from aDNA genomes, that is consistent across libraries with different sequencing strategies and read depths, we developed the `impHet` pipeline. `impHet` uses imputed genotypes from GLIMPSE2 [44, 47] to compute heterozygosity estimates. Current approaches to generating heterozygosity estimates from imputed genotypes rely on stringent filtering by minor allele frequency (MAF) to remove variants with low imputation confidence [42]. However, heterozygosity estimates computed from subsets of variants with high MAF values are expected to be inflated (Fig. S1). Rather than filtering out variants with low confidence, we determine the expected accuracy of heterozygosity estimates in relation to the coverage of the input genome. This allows us to filter input genomes using a coverage threshold that ensures a certain level of accuracy, and avoids biasing the heterozygosity estimate by excluding variants with low MAF values.

`impHet` handles the preparation of files for GLIMPSE2 and the different stages of imputation, and is designed to process many genomes in parallel. The output of `impHet` includes heterozygosity estimates for the genomic regions provided by the user, for each genome, as well as *β*_*H*_ values and selection coefficient estimates for each region. Further details on the implementation and instructions for using the pipeline are available at https://github.com/Greenbaum-Lab/imphet.git.

To process imputed genotypes to heterozygosity values, `impHet` filters the genotypes from GLIMPSE2 using `bcftools +setGT`. When processing multiple genomes, this is performed on a merged BCF file containing all genomes. `impHet` allows users to define the threshold of genotype probability for imputed genotypes; here we used a threshold of *GP* ≥ 0.9 for all analyses. Genotypes below the threshold are assigned as missing. The filtered BCF is then converted to `PLINK BED` format using `PLINK 1.90b` [68]. All downstream analyses are conducted on the `PLINK BED` file using the Python package `bed reader` [69]. To generate heterozygosity estimates for specific genomic regions, we compute for each genome the proportion of heterozygous variants in the region from the number of variants with non-missing genotypes. These values are then used to compute *β*_*H*_ values, as detailed in the ‘*Genomic scan for balancing selection using β*_*H*_ ‘ section below.

### 4.3 Validation of heterozygosity estimates

GLIMPSE2 provides a pipeline for testing the average concordance of variants in relation to their MAF in the reference panel [44, 47]. This result can be used as an indication of the appropriate MAF threshold for filtering imputed genotypes generated by GLIMPSE2. However, there is no equivalent pipeline for assessing the accuracy of per-genome estimates generated by aggregating imputed genotypes, such as heterozygosity. In addition, downsampling procedures often subsample reads uniformly [44]. This is suitable for comparing shotgun sequenced genomes with different depths; however, uniform subsampling cannot mimick SNP capture sequencing. This distinction is important, because SNP capture sequencing can achieve high accuracy imputation at target regions of the SNP capture panel, and low accuracy imputation at non-target regions. On the other hand, generating simulated low-coverage genomes by subsampling reads only at the target regions of the SNP capture panel ignores the effect of off-target reads, which can be substantial for imputation [45].

To address these issues, we designed a downsampling scheme that recapitulates read distribution patterns of different low-coverage genomes for a high-coverage genome (Fig. 1B). As opposed to uniform subsampling, this method generates meaningful replicates for the same high-coverage genome: here each replicate contains the same data as the original, high-coverage genome, but “as if” it was sequenced with the sequencing method and depth of the low-coverage genome.

Read depth is first binned into windows of 50bp in the low- and high-coverage genome BAM files using `bedtools`. To generate a downsampled file, for each window of 50bp with depth ≥ 1 in the high-coverage genome, reads are sampled from the high-coverage genome with the probability min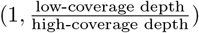, according to the bin of the starting position of the read. A bin size of 50bp was used as it is similar to the average length of aDNA reads, and after testing the effect of different bin sizes on the similarity of the coverage and average depth of the low-coverage genome and the downsampled output.

To measure the accuracy of the estimate, we selected 8 high-coverage genomes with different levels of UDG treatment (`SI table1.xlsx`). Each high-coverage genome was downsampled using 1000 low-coverage genomes, include 50% genomes generated by SNP capture and 50% genomes generated by shotgun sequencing. The data were then processed by `impHet` to produce estimated heterozygosity values for windowed regions of the genome. Heterozygosity values were also computed using the non-imputed genotypes, to compare to the values from `impHet`. In addition, heterozygosity values were computed for the full high-coverage genomes, both with and without imputation (Fig. 1C).

### 4.4 Genomic scan for balancing selection using *β*_*H*_

To develop a genomic scan for balancing selection, we introduce a simple statistic, *β*_*H*_, which measures the change in heterozygosity per 1000 years. *β*_*H*_ is computed as *β*_*H*_ = *β*_1_ ∗ 1000, where *β*_1_ the slope of the linear regression of heterozygosity over time. We use the *p*-value of the regression to determine the significance of the *β*_*H*_ value, using the Benjamini-Hochberg procedure with *α* = 0.01. To determine the stability of the statistic, we bootstrap the genomes in each group by resampling with replacement the genomes with 1000 repeats. We retain hits that are significant in the regression for the full group and at least 95% of bootstrap replicates.

We ran a genomic scan using two approaches: using annotated genes (GENCODE v19), and using 10Kb genomic windows. This window size was chosen based on the size of significant genes, and the loss of sensitivity and resolution at large window sizes. To assess the effect of our choice of annotation, we also ran the genomic scan with the GENCODE v47 annotation, lifted to the hg19 assembly. We filtered out genes shorter than 500bp, and selected genes with a “KNOWN” gene status for GENCODE v19, and genes with a HGNC ID for GENCODE v47. For MHC class boundaries, we followed [70], lifted to hg19 GENCODE v19 positions: 29690552–31478901 for MHC class I (HLA-F to MICB), 32374958–32374958 for MHC class II (the end of BTNL2 to HLA-DPA3), and 31486754–32374958 for MHC class III (PPIAP9 to BTNL2).

### 4.5 Accounting for ancestry differences in genomic scans of all genomes

For the Western and Eastern groups, we selected samples based on a number of criteria, including ancestry similarity (Fig. S4). This reduces the effect of ancestry differences among genomes in the scan on our results. To run a genomic scan using all 3846 genomes in the AADR that passed our coverage threshold (top panel in Fig. 2C), we used as covariates the first two ancestry components of an ADMIXTURE model of ancient genomes in the AADR (*K* = 3) and the genome-wide heterozygosity estimate. We first fit a multiple regression using the three covariates for each genomic region, and then used the residuals of the heterozygosity of each genomic region for the scan. We used the same procedure for the genomic scan of all 3846 genomes excluding the Western and Eastern groups (middle panel in Fig. 2C).

### 4.6 Estimating linkage disequilibrium between genes

Because some of the significant genes from the genomic scans are clustered along the genome, we assessed the average LD between variants in all pairs of genes situated less than 100Kb apart. For this analysis we used the Southern Europe LD panel from gnomAD v2 [71]. We first selected variants from the LD panel that are situated in each of the significant genes in the genomic scan. We then computed *r*^2^ values for each pairs of variants between the two genes and average them to generate a single value for each gene pair,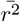. For variants overlapping between genes, we assumed a value of *r*^2^ = 1. To identify clusters of genes with substantial LD or overlap, we used the 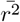 matrix to isolate connected components (edges with 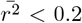 or no overlap between genes were removed).

### 4.7 Temporal trends of heterozygosity

To track temporal trends of heterozygosity in the MHC and whole genome, we analyzed the mean heterozygosity of genomes using a 2000-year sliding window with a step size of 500 years, in each group. We excluded windows with less than 10 genomes. We used bootstrapping to generate confidence intervals of heterozygosity in each window, resampling the genomes in each window with 1000 repeats.

To identify periods during which heterozygosity increased substantially, we computed *β*_*H*_ values for a 3000-year sliding window, with a step size of 500 years, in each group. The larger window is necessary to capture a change in heterozygosity (as opposed to the 2000-year window, the purpose of which is only to estimate the heterozygosity at a specific period). We removed windows with a midpoint earlier than 9000BP due to noise in the signal during this period (no significant signal was identified earlier than 9000BP).

### 4.8 Estimating effective balancing selection coefficients

In order to broadly estimate and compare the level of balancing selection in temporal windows we used a simplified mathematical model. We first consider heterozygote advantage (HA) as a mechanism for balancing selection, and use a symmetric over-dominance model. To simplify the multi-allelic nature of the MHC region into a tractable mathematical model, for each MHC region we calculated an *effective allele frequency* (*p*_*e*_) from the observed mean heterozygosity, assuming Hardy-Weinberg equilibrium. In other words, if 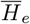 is the mean heterozygosity in a given MHC region, we solve

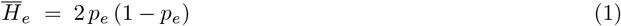

for *p*_*e*_ (taking the minor allele frequency, for consistency). This conversion lets us treat the region as if it were a single biallelic locus with heterozygosity 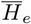. While this single-frequency approach necessarily omits the complexities of large MHC regions, it provides a consistent baseline for comparing balancing selection levels in different populations and across different time points.

To compute the effective selection coefficient between two time points, we compare the changes in the effective allele frequencies under the HA model. Here, assuming that the fitness of the heterozygote is 1, and the fitness of the two homozygote genotypes is 1 − *s*, then the change of allele frequency over a single generation can be formulated as:

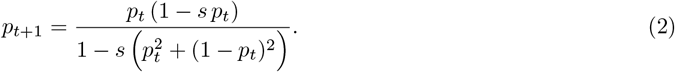

Given effective allele frequencies *p*_0_ and *p*_1_ computed in two time windows separated by *T* generations (we assumed generation time of 25 years; [72]), we used a bisection search on the *s* values to identify an effective selection coefficient *s* that yields a change from *p*_0_ to *p*_1_ in *T* iterations of Eq. 2.

Another mechanism that can generate balancing selection is negative-frequency dependent selection (NFDS). If we use a simple biallelic haploid model where the fitness of an allele with frequency *p*_*t*_ at time *t* is given by 1 − *sp*_*t*_ [73], then the change in frequency can also be described by Eq. 2. Therefore, the procedure described here for quantifying balancing selection is based on a simplistic model (symmetric heterozygote advantage or linear haploid negative-frequency dependence), but is not specific to a particular evolutionary mechanism.

In order to provide a null distribution to which we can compare the estimated selection coefficients for MHC classes, we used genome-wide heterozygosity change to generate a distribution of *s* values. For this, we computed selection coefficients for all regions of 1Mb (2729 windows excluding the MHC). This window size was chosen as it is of the same scale as MHC classes.

### 4.9 Gradients of signals of selection

To represent a color gradient on a timeline for the signals of selection we identified in MHC classes I and II (Fig. 3E–F), we used the values from our analysis of temporal trends of *β*_*H*_ (Fig. 3C–D). Values below *β*_*H*_ = 0.004 were set to 0, and remaining values were normalized using the maximum value of *β*_*H*_ across all MHC classes.

### 4.10 Climate timelines

To generate estimates of climate differences over time in each group, we used the CHELSA-TraCE21k dataset [74], which provides 100-year interval estimates at approximately 1km geographic resolution of different climate measures for the past 21,000 years. To summarize the measures, we conducted a principle component analysis (PCA) of 19 climatic parameters, in 1,000-year intervals at 100km resolution (generated by averaging 1km resolution values at 1000-year intervals). The first principal component reflects mainly changes in temperature, while the second reflects changes in precipitation. To summarize these components for each group, we averaged the values contained in the convex hull of the geolocation of individuals in each group (for the entire 12,000-year period). Because each group has a different range of values for each principal component, we normalized the timeline values relative to those in the earliest time window.

### 4.11 Correlation of genomic scan between groups

To compare the results of the genomic scans for the Western and Eastern groups, we compute a linear regression for the *β*_*H*_ values from each group, for genes or windows that were significant in either group. We removed from this analysis genes that overlap other genes with at least 50% of their length (the shorter gene is removed).

### 4.12 Temporal trends of heterozygosity for significant genes

To compare the temporal trends of genes identified in our genomic scan to the temporal trends of whole MHC classes, we analyzed the mean observed heterozygosity in each group using a 2000-year sliding window. To differentiate increases in observed heterozygosity caused by increased mixing within the population, as opposed to increased genetic diversity across the group, we also analyzed the expected heterozygosity using the same sliding window.

## Acknowledgments

We thank David Gokhman, Liran Carmel and Shai Carmi for input at various stages of the study’s development, and Adriana Morales-Guerrero and Simon Fishilevich for assistance with data curation. We would also like to thank members of the Greenbaum, Slon, Carmel and Gokhman labs for feedback. KDH was supported by a fellowship from the Council for Higher Education of Israel and by the Minerva Center for the study of Population Fragmentation (MCPF). VS was supported by the Israel Science Foundation (grant number 2075/22).

## 5 Author contributions

The study was conceived by GG and VS The study was designed by GG and KDH, with input from NM, YJ and VS. Software was written by KDH, YT and MBY. Analyses were conducted by KDH, MBY and YT. The original draft was written by KDH and GG. NM, YJ and VS contributed additional writing. The study was supervised by GG and KDH. All authors reviewed the final manuscript.

## 6 Competing interesting

The authors declare no competing interests.

## 7 Data and code accessibility

All source files used are publicly available and accession numbers are provided in SI table1.xlsx. The `impHet` source code is available at https://github.com/Greenbaum-Lab/imphet.git. Additional code used to process the data is available as a Jupyter notebook at https://github.com/Greenbaum-Lab/imphet_figures.git. Heterozygosity estimates are available to visualize through the DORA platform (https://dora.modelrxiv.org).

