## Supplemental Information for "Genomic signatures of increasing disease burden in recent prehistory"

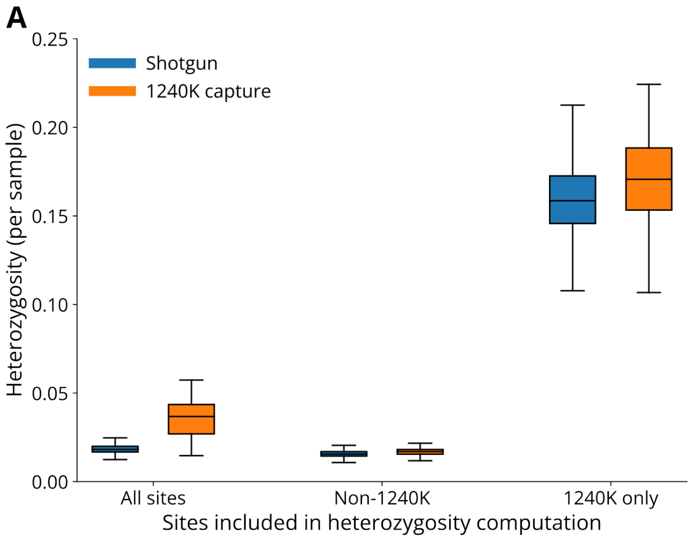


Figure S1: **Biased heterozygosity estimates from SNP capture panels**. Shown are measured heterozygosity values for shotgun (blue) or 1240K SNP capture (orange) read distributions of the Yana1 genome, computed using different subsets of SNPs. Read distributions were generated by resampling reads from the original Yana1 high-coverage genome using our downsampling pipeline for 1000 low-coverage genomes (50% generated by shotgun sequencing, 50% by 1240K SNP capture), filtered for coverage (number of variants with a read depth of at least 4) of at least 100 variants in chromosome 6 ($n = 997$).

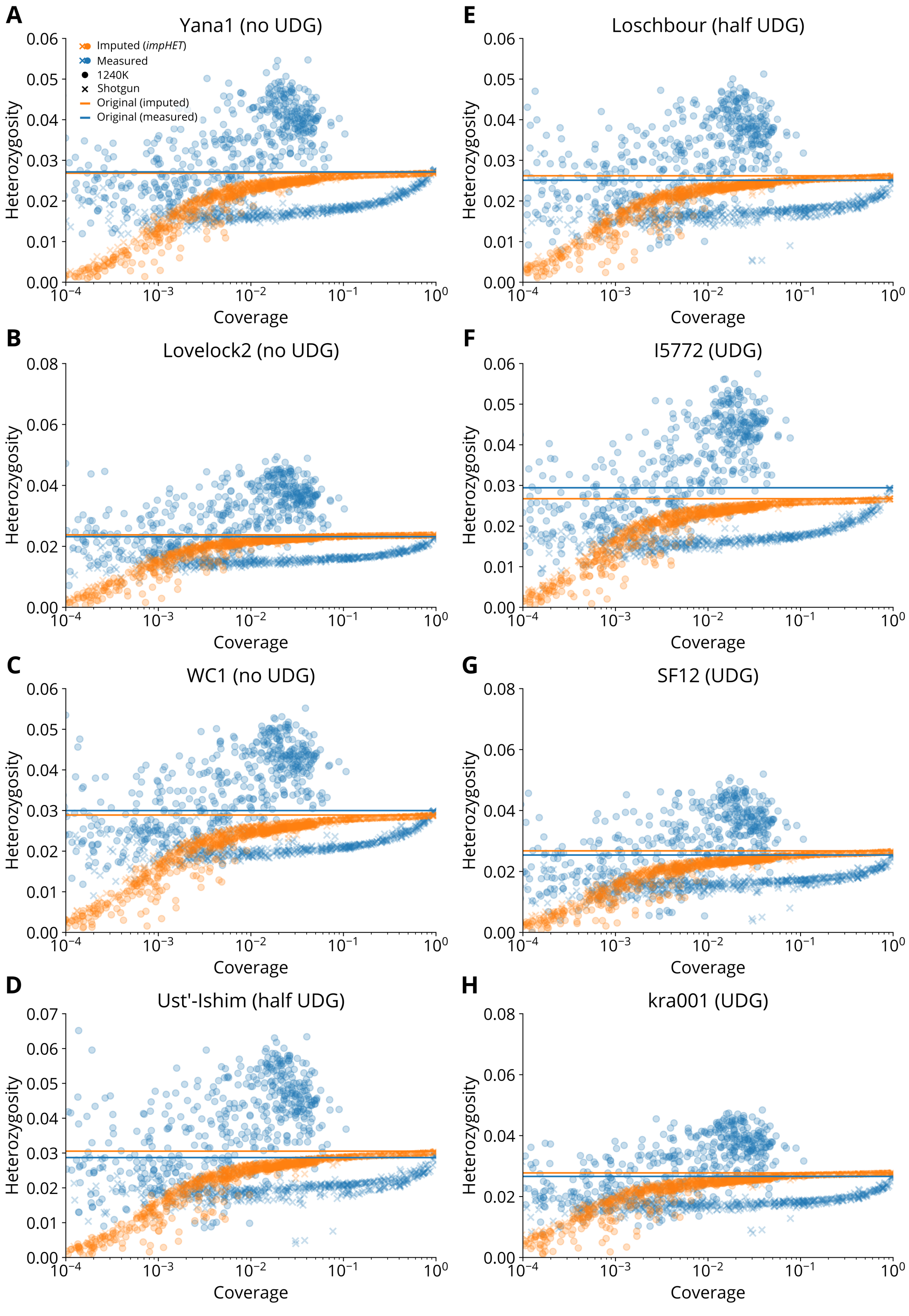


Figure S2: **Accuracy of `impHet` heterozygosity estimate in chromosome 6 (excluding the MHC region)**. (A–H) Heterozygosity estimates either non-imputed (blue circles and crosses) or from `impHet` (orange circles and crosses) for genomes generated by downsampling high-coverage genomes with different UDG treatments. 1000 low-coverage read-distribution templates were used for each panel. Coverage relates to the proportion of variants with a read depth $\geq 4$ in each downsampled genome, out of all variants in the panel for the specific genomic region. Horizontal lines indicate the "ground truth" heterozygosity for the full coverage of the source genome, either non-imputed (blue) or from `impHet` (orange). For a list of genomes use, see `SI_table1.xlsx`.

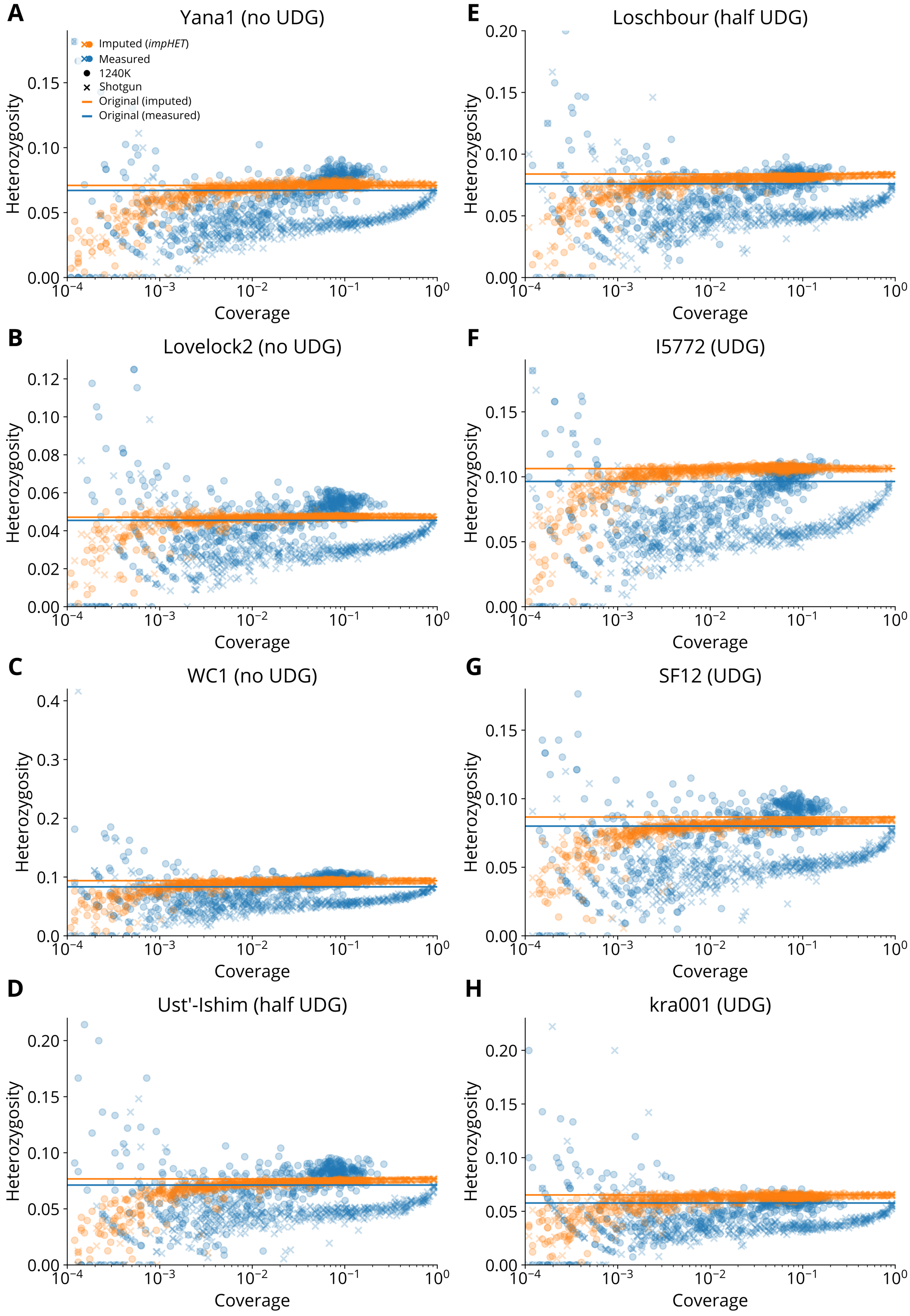


Figure S3: **Accuracy of `impHet` heterozygosity estimate in the MHC region**. (A–H) Heterozygosity estimates either non-imputed (blue circles and crosses) or from `impHet` (orange circles and crosses) for genomes generated by downsampling high-coverage genomes with different UDG treatments. 1000 low-coverage read-distribution templates were used for each panel. Coverage relates to the proportion of variants with a read depth $\geq 4$ in each downsampled genome, out of all variants in the panel for the specific genomic region. Horizontal lines indicate the "ground truth" heterozygosity for the full coverage of the source genome, either non-imputed (blue) or from `impHet` (orange). For a list of genomes use, see `SI_table1.xlsx`.

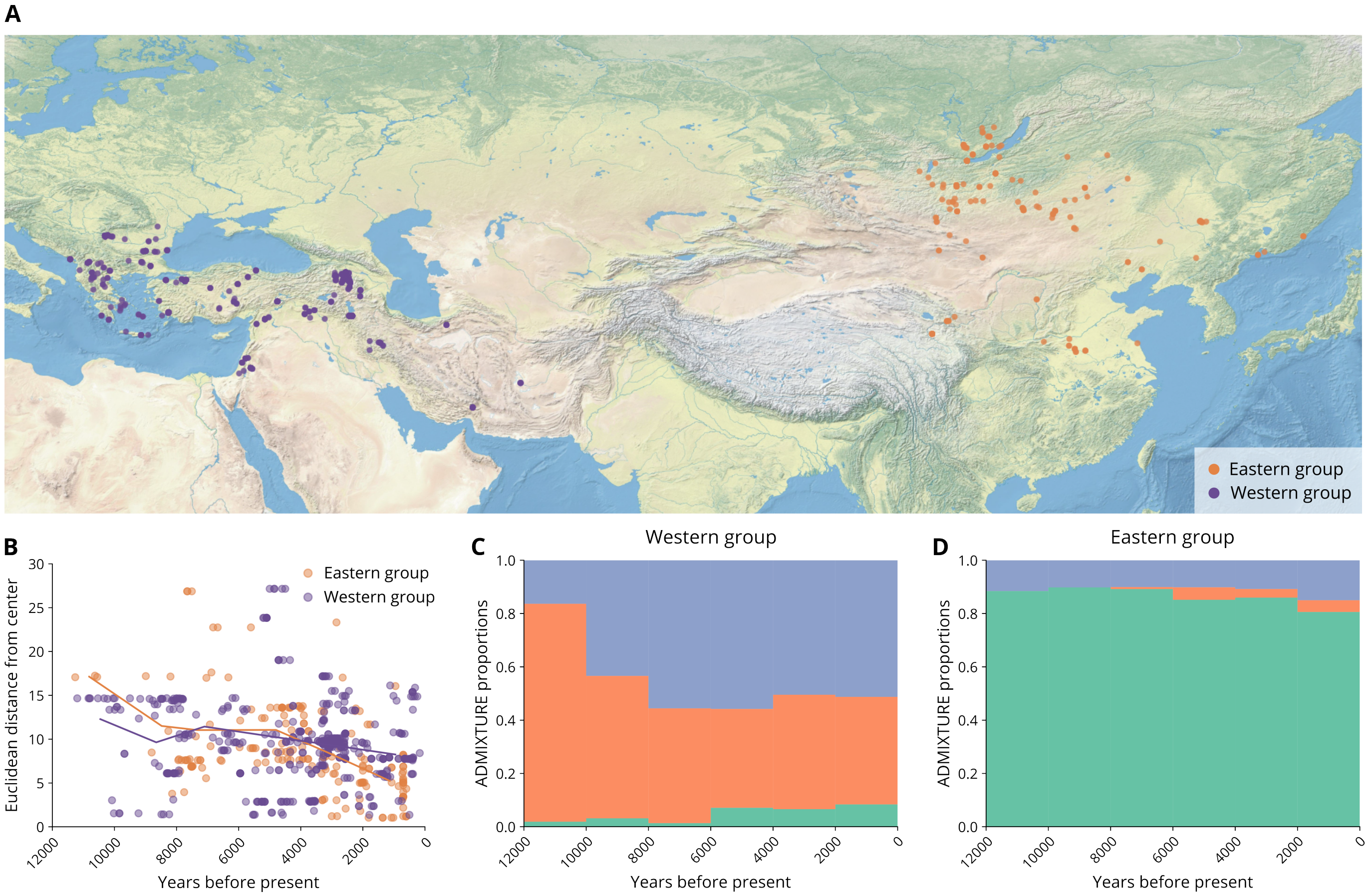


Figure S4: **Geographical locations and ancestry of individuals used for each group**. (A) Geographical distribution of the Western group (purple circles) and Eastern group (orange circles). (B) Euclidean distance of each individual in the Western group (purple circles) and Eastern group (orange circles) from the from the centroid (mean geographic coordinates) of its group. Solid lines show the mean distance of the Western group (purple) and Eastern group (orange). (C–D) ADMIXTURE result ($K = 3$) for genomes in the Western group (C) and Eastern group (D), binned in 2000-year windows, showing the change in ancestry components over time.

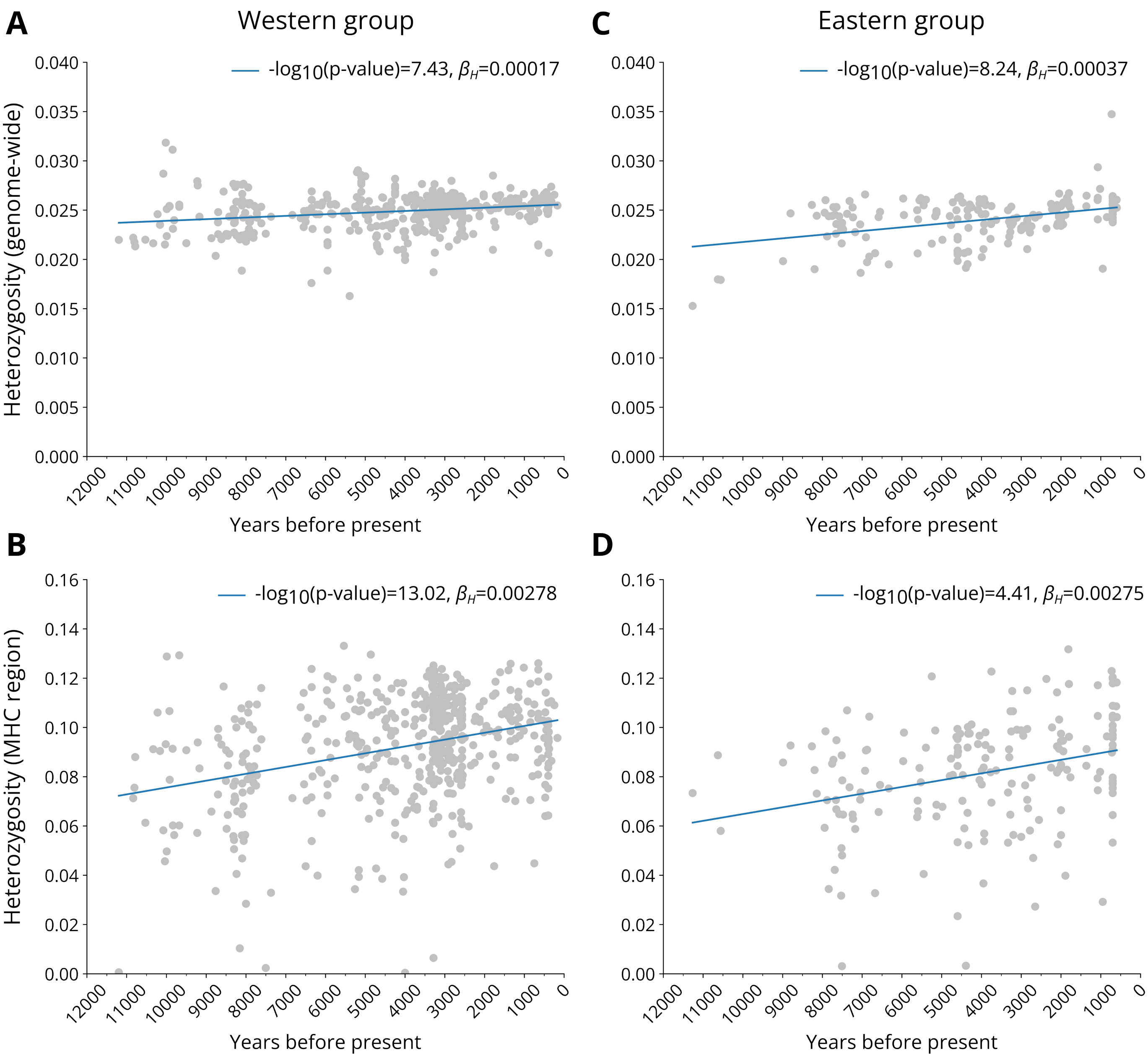


Figure S5: **Linear regressions of date (years before present) and heterozygosity.** Circles represent individual genomes. The solid blue line indicates the slope of the linear regression. The p-value and $\beta_H$ are shown in the legend of each panel. (A–B) Linear regression of heterozygosity and date for the Western group, genome-wide (A) and in the MHC (as shown in Figure 2 in the main text) (B). (C–D) Linear regression of heterozygosity and date for the Western group, genome-wide (C) and in the MHC (D).

| Gene | Genomic position | $\beta_H$ (bootstrap %) | MHC | Known immune function | Previous findings |
|---|---|---|---|---|---|
| CCR3 | 3:46205096-46308197 | 0.00245 (99.5%) | - | [1, 2] | [3] |
| COL4A3BP | 5:74664311-74807963 | 0.00105 (96.6%) | - | No | N/A |
| POLK | 5:74807581-74896969 | 0.00108 (95.5%) | - | No | N/A |
| P4HA2 | 5:131527531-131631008 | 0.00185 (96.7%) | - | No | N/A |
| GPX3 | 5:150400124-150408554 | 0.00318 (96.6%) | - | No | N/A |
| HCG4P11 | 6:29690758-29691748 | 0.00822 (99.6%) | I | No | N/A |
| HCG4B | 6:29893760-29894750 | 0.00124 (98.0%) | I | No | N/A |
| HLA-A | 6:29909037-29913661 | 0.01026 (100.0%) | I | [4, 5] | [6] |
| HCG4P5 | 6:29909852-29910844 | 0.01330 (100.0%) | I | No | N/A |
| DHFRP2 | 6:31334129-31334675 | 0.00994 (100.0%) | I | No | N/A |
| XXbac-BPG181B23.6 | 6:31430505-31431113 | 0.00513 (98.6%) | I | No | N/A |
| PRRC2A | 6:31588497-31605548 | 0.00211 (99.6%) | III | No | N/A |
| BAG6 | 6:31606805-31620482 | 0.00212 (98.6%) | III | [7] | N/A |
| APOM | 6:31620193-31625987 | 0.00145 (99.1%) | III | [8] | N/A |
| LY6G5B | 6:31637944-31641553 | 0.00233 (100.0%) | III | No | N/A |
| CLIC1 | 6:31698358-31707540 | 0.00091 (100.0%) | III | [9] | N/A |
| HSPA1L | 6:31777396-31783437 | 0.00133 (96.1%) | III | No | N/A |
| HSPA1B | 6:31795512-31798031 | 0.00216 (99.7%) | III | No | N/A |
| EHMT2 | 6:31847536-31865464 | 0.00102 (97.4%) | III | No | N/A |
| DXO | 6:31937587-31940069 | 0.00120 (95.0%) | III | No | N/A |
| CYP21A1P | 6:31973413-31976228 | 0.00120 (98.9%) | III | No | N/A |
| C4B | 6:31982539-32003195 | 0.00268 (99.6%) | III | [10] | N/A |
| CYP21A2 | 6:32006042-32009447 | 0.00223 (99.6%) | III | No | N/A |
| TNXB | 6:32008931-32083111 | 0.00137 (99.4%) | III | No | N/A |
| ATF6B | 6:32065953-32096030 | 0.00107 (99.4%) | III | [11] | N/A |
| FKBPL | 6:32096484-32098068 | 0.00418 (99.8%) | III | No | N/A |
| PRRT1 | 6:32116136-32122150 | 0.00102 (97.1%) | III | No | N/A |
| PPT2 | 6:32121218-32134011 | 0.00202 (100.0%) | III | No | N/A |
| EGFL8 | 6:32132360-32136058 | 0.00214 (99.3%) | III | No | N/A |
| AGPAT1 | 6:32135989-32145873 | 0.00173 (99.6%) | III | No | N/A |
| RNF5 | 6:32146131-32151930 | 0.00286 (100.0%) | III | [12] | N/A |
| AGER | 6:32148745-32152101 | 0.00249 (100.0%) | III | [13] | [14] |
| PBX2 | 6:32152512-32157963 | 0.00249 (100.0%) | III | No | N/A |
| GPSM3 | 6:32158543-32163300 | 0.00173 (99.0%) | III | [15] | N/A |
| HLA-DRB6 | 6:32520490-32527799 | 0.00987 (98.7%) | II | [16] | N/A |
| HLA-DQB1 | 6:32627244-32636160 | 0.01273 (95.3%) | II | [17, 18] | [19-21] |
| AC011290.5 | 7:39608975-39609797 | 0.00624 (95.7%) | - | No | N/A |
| LHPP | 10:126150403-126306457 | 0.00167 (98.3%) | - | No | N/A |
| OAS1 | 12:113344582-113369990 | 0.00536 (95.5%) | - | [22] | [23] |
| HSD17B1 | 17:40701232-40707231 | 0.00269 (97.5%) | - | No | N/A |
| COASY | 17:40713485-40718295 | 0.00219 (95.2%) | - | No | N/A |
| FAM134C | 17:40731531-40762641 | 0.00194 (99.1%) | - | No | N/A |
| TUBG1 | 17:40761694-40767252 | 0.00080 (95.2%) | - | No | N/A |
| TUBG2 | 17:40811323-40819024 | 0.00178 (96.6%) | - | No | N/A |
| CCR10 | 17:40830907-40835935 | 0.00280 (97.0%) | - | [24, 25] | N/A |
| CNTNAP1 | 17:40834631-40851832 | 0.00200 (100.0%) | - | No | N/A |
| GRB2 | 17:73314157-73401790 | 0.00044 (99.9%) | - | No | N/A |

Table S1: **List of significant genes with positive $\beta_H$ values in the Western group genomic scan**. Shown are the annotation details of each gene (chromosome and position), $\beta_H$ value, and the percentage of bootstrap repeats in which it was significant. If relevant, the MHC class of the gene is shown. Citations are provided for genes with known immune functions, or that appeared as hits in previous genomic scans for selection.

| Gene | Genomic position | $\beta_H$ (bootstrap %) | MHC | Known immune function | Previous findings |
|---|---|---|---|---|---|
| PDE4DIP | 1:144836157-145076186 | -0.00046 (99.4%) | - | No | N/A |
| SEC22B | 1:145096220-145116922 | -0.00103 (98.3%) | - | No | N/A |
| RP11-433J22.3 | 1:147249700-147261065 | -0.00254 (96.8%) | - | No | N/A |
| KCNT2 | 1:196194909-196578355 | -0.00085 (98.7%) | - | No | N/A |
| VTI1BP1 | 3:99944218-99944917 | -0.00350 (96.7%) | - | No | N/A |
| RPL9P15 | 3:154394377-154394908 | -0.00158 (95.8%) | - | No | N/A |
| DUSP22 | 6:291630-351355 | -0.00014 (95.4%) | - | No | N/A |
| HOXA3 | 7:27145803-27192200 | -0.00143 (99.3%) | - | [26] | N/A |
| HOXA-AS2 | 7:27147396-27173921 | -0.00148 (99.3%) | - | [27] | N/A |
| HOXA-AS3 | 7:27169596-27195542 | -0.00116 (98.2%) | - | [28] | N/A |
| RP11-351I21.6 | 8:12236188-12237774 | -0.00071 (98.8%) | - | No | N/A |
| RP11-38L15.3 | 10:46951472-46966835 | -0.00032 (95.5%) | - | No | N/A |
| RP11-508M1.7 | 10:48988121-48998766 | -0.00111 (96.6%) | - | No | N/A |
| RP11-579D7.8 | 12:49194359-49194869 | -0.00173 (97.5%) | - | No | N/A |
| RP13-395E19.2 | 15:32605327-32607855 | -0.00049 (95.0%) | - | No | N/A |
| DUOX2 | 15:45384848-45406542 | -0.00133 (99.8%) | - | [29] | [21, 30] |
| SLC24A5 | 15:48413169-48434869 | -0.00072 (99.7%) | - | No | [19, 31] |
| MYEF2 | 15:48431625-48470714 | -0.00064 (99.7%) | - | No | N/A |
| KIAA0895L | 16:67209505-67217943 | -0.00071 (95.8%) | - | No | N/A |
| LRRC29 | 16:67241042-67260951 | -0.00092 (98.4%) | - | No | N/A |
| AC040160.1 | 16:67244251-67260930 | -0.00077 (98.0%) | - | No | N/A |
| KCNJ12 | 17:21279509-21323179 | -0.00076 (98.0%) | - | No | N/A |
| RIMBP3 | 22:20456003-20461786 | -0.00031 (97.9%) | - | No | N/A |
| SHISA8 | 22:42307297-42310570 | -0.00379 (95.3%) | - | No | N/A |

Table S2: **List of significant genes with negative $\beta_H$ values in the Western group genomic scan**. Shown are the annotation details of each gene (chromosome and position), $\beta_H$ value, and the percentage of bootstrap repeats in which it was significant. If relevant, the MHC class of the gene is shown. Citations are provided for genes with known immune functions, or that appeared as hits in previous genomic scans for selection.

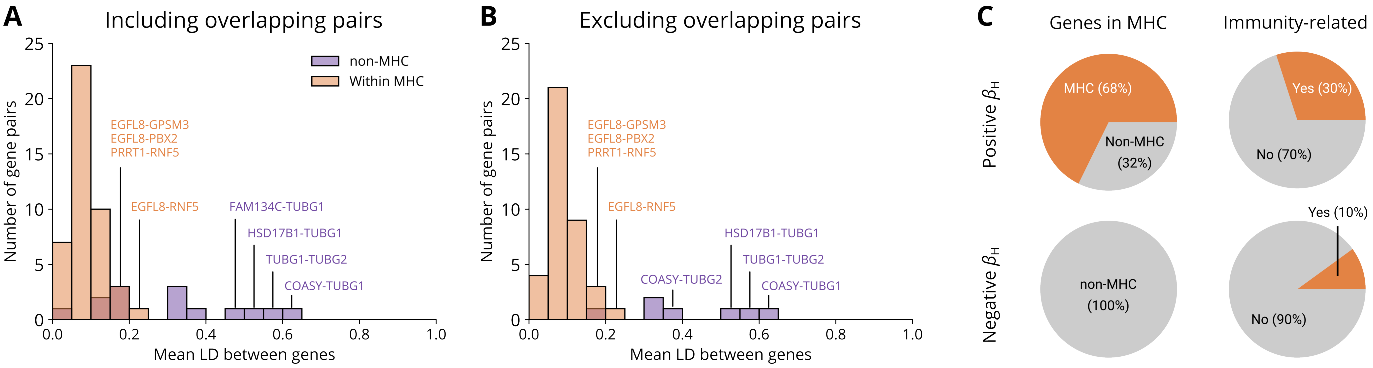


Figure S6: **Linkage disequilibrium ($\bar{r^2}$) in gene pairs for significant genes identified in the genomic scan for the Western group**. (A) Distribution of $\bar{r^2}$ values for pairs of significant genes within 100Kb of each other. Values for gene pairs were computed using $r$ values for variant pairs between the two genes for which data was available in the gnomAD v2 Southern Europe LD panel [32]. For overlapping genes, overlapping variants present in the LD panel were assigned an $r$ value of 1. Shown are the $\bar{r^2}$ values for all gene pairs within the MHC or outside the MHC (there were no pairs between MHC classes, or between the genes in the MHC and genes outside the MHC). The gene pairs with the four highest $\bar{r^2}$ values are annotated for pairs in the MHC (orange) and outside the MHC (purple). (B) Distribution of $\bar{r^2}$ values for pairs of significant genes within 100Kb of each other, excluding pairs of overlapping genes. The gene pairs with the four highest $\bar{r^2}$ values are annotated for pairs in the MHC (orange) and outside the MHC (purple). (C) Pie charts showing enrichment of MHC or immune-related genes (outside the MHC) in significant hits of the genomic scan in Figure 2 in the main text, after removing genes within 100Kb of each other with $\bar{r^2} \geq 0.2$. The results are separated to positive (increase in heterozygosity) and negative (decrease in heterozygosity) trends.

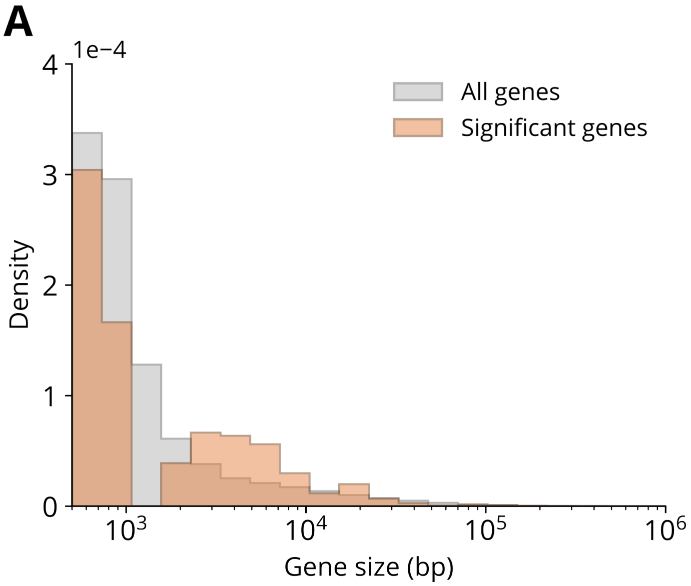


Figure S7: **The effect of gene length on discovery in $\beta_H$ genomic scan**. Plotted are histograms of gene length in base pairs (bp) for significant genes (orange) and all genes with "KNOWN" status in the GENCODE v19 annotation (gray). Only genes at least 500bp in length are included.

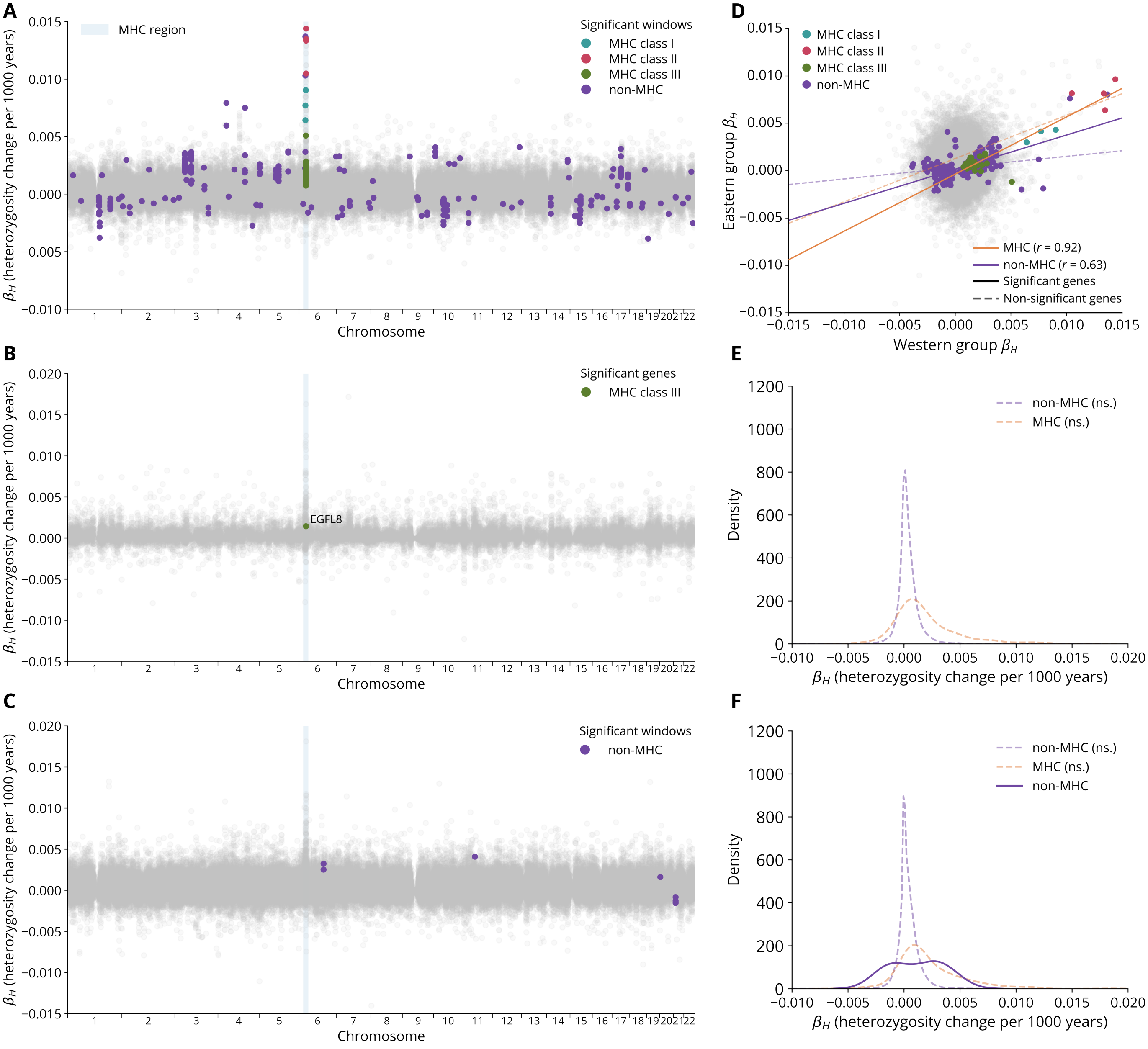


Figure S8: **Genome-wide scan using $\beta_H$ in genes and fixed-size genomic windows**. (A) Values of $\beta_H$ for fixed-size windows of 10Kb in the Western group. The MHC region is highlighted in blue. Significant windows are colored turquoise for MHC class I, magenta for MHC class II, green for MHC class III and purple for all other genes. Non-significant windows are colored gray. (B) Values of $\beta_H$ for annotated genes in the Eastern group. The MHC is highlighted in blue. The single significant gene, which is in MHC class III, is colored green. Non-significant genes are colored gray. (C) Values of $\beta_H$ for fixed-size windows of 10Kb in the Eastern group. The MHC is highlighted in blue. Significant windows outside the MHC are colored purple (there are no significant windows in the MHC). Non-significant genes are colored gray. (D) Correlation between the $\beta_H$ values of significant 10Kb windows between the Western group and Eastern group. Windows with significant results in either group are colored according to the genomic region: turquoise for MHC class I, magenta for MHC class II, green for MHC class III and purple for all other genes. Windows with no significant result are colored gray. The solid orange line is the regression result for windows in the MHC with a significant regression, and the dashed orange line is the result for windows in the MHC with no significant regressions. The solid purple line is the result for windows outside the MHC with a significant regression, and the dashed purple line is the result for windows outside the MHC with no significant regressions. (E) Kernel density estimates for $\beta_H$ values of genes in the Eastern group (the genome-wide distribution of which appears in B). Shown are kernel density estimates for non-significant genes in MHC and non-MHC, represented by orange and purple dashed lines, respectively. (F) Kernel density estimates for $\beta_H$ values from a genomic scan using fixed-size windows of 10Kb in the Eastern group (the genome-wide distribution of which appears in C). Shown are significant non-MHC genes (solid purple line), with non-significant genes in MHC and non-MHC represented by orange and purple dashed lines, respectively.

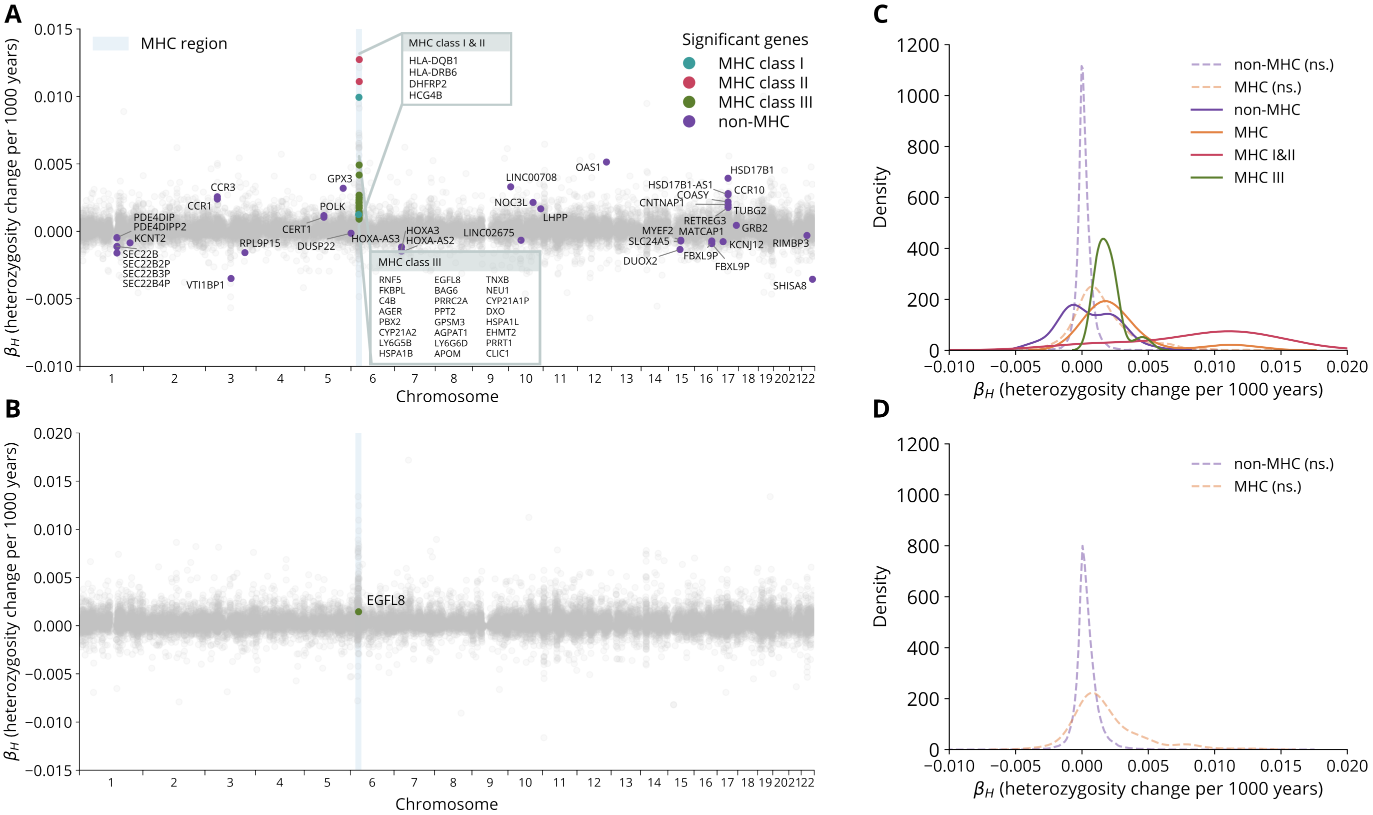


Figure S9: **Genome-wide scan using $\beta_H$ using the GENCODE v47 annotation**. (A) Values of $\beta_H$ for annotated genes in the Western group. The MHC is highlighted in blue. Significant genes are colored turquoise for MHC class I, magenta for MHC class II, green for MHC class III and purple for all other genes. Non-significant genes are colored gray. All significant genes are annotated, with boxes for the densely clustered MHC genes. (B) Values of $\beta_H$ for annotated genes in the Eastern group. The MHC is highlighted in blue. The single significant gene, which is in MHC class III, is colored green. Non-significant genes are colored gray. (C) Kernel density estimates for $\beta_H$ in the Western group, the genome-wide distribution of which is shown in (A). Shown are significant MHC genes (orange), MHC class I&II genes (magenta), MHC class III genes (green) and non-MHC genes (purple). Kernel density estimates for non-significant genes in MHC and non-MHC are represented by orange and purple dashed lines, respectively. (D) Kernel density estimates for $\beta_H$ in the Eastern group, the genome-wide distribution of which is shown in (B). Shown are kernel density estimates for non-significant genes in MHC and non-MHC, represented by orange and purple dashed lines, respectively.

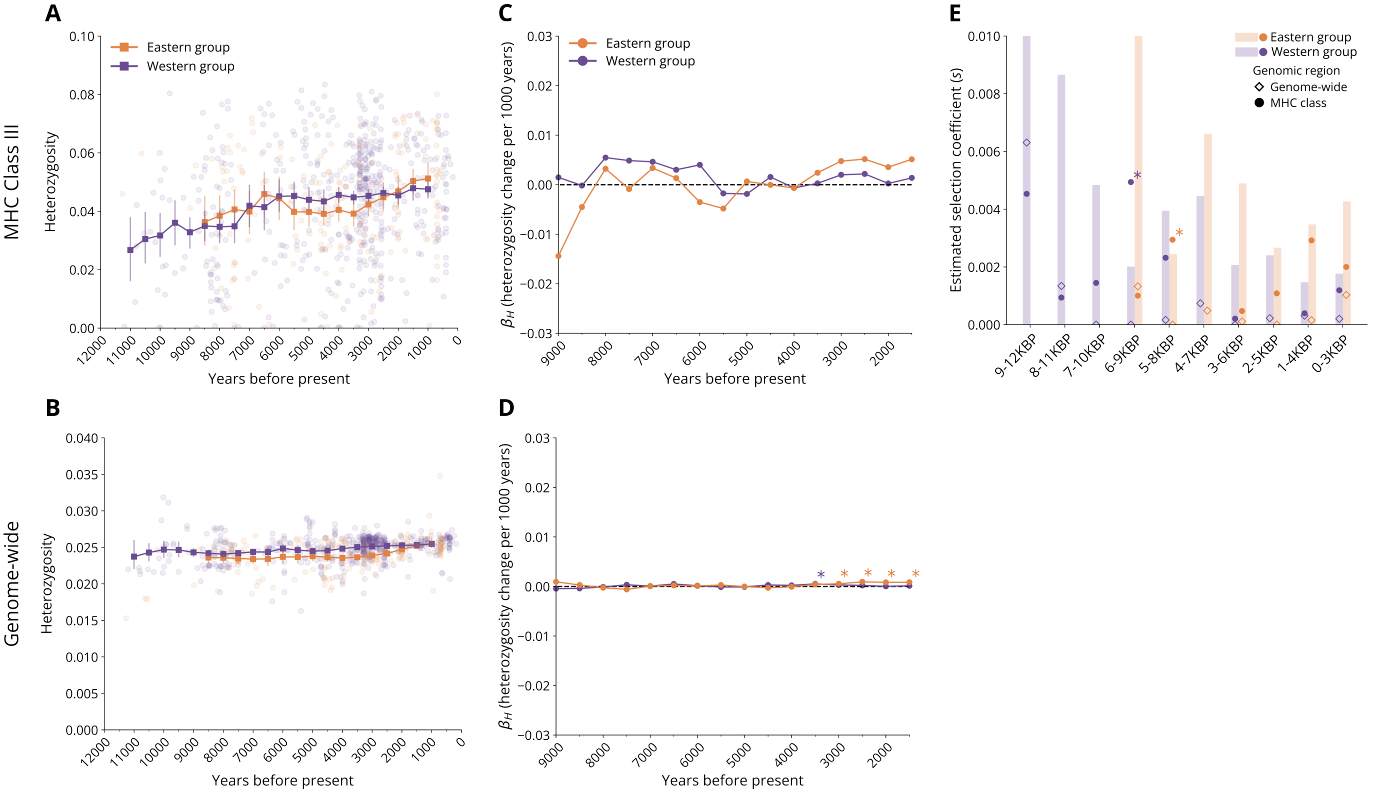


Figure S10: **Temporal trends of heterozygosity in MHC class III and genome-wide**. (A–B) Sliding window analysis of heterozygosity of MHC class III (A) and the entire genome (B), in the Western group (purple) and Eastern group (orange). Solid lines with squares represent heterozygosity calculated over 2000-year windows centered around each midpoint on the x-axis. Circles represent heterozygosity values of individual genomes according to their date (years before present). Error bars indicate the 95% confidence intervals of the mean, generated using bootstrapping. Windows with less than 10 genomes are not shown. (C–D) Temporal analysis of $\beta_H$ in MHC class III (C) and the entire genome (D) in the Western group (purple) and Eastern group (orange). Solid lines with circles represent $\beta_H$ values calculated over 3000-year windows centered around each midpoint on the x-axis. Asterisks indicate significant regressions for the specific window. (E) Estimates of balancing selection in MHC class III in the Western group (purple) and Eastern group (orange). Each column represents an overlapping 3000-year period. Shaded bars represent the extent of the upper 95% confidence interval of the genome-wide signal. Empty diamonds represent the genome-wide signal, and filled circles represent the signal of the specific MHC class. Windows in which the MHC class signal is outside the confidence interval of the genome-wide signal are indicated with asterisks.

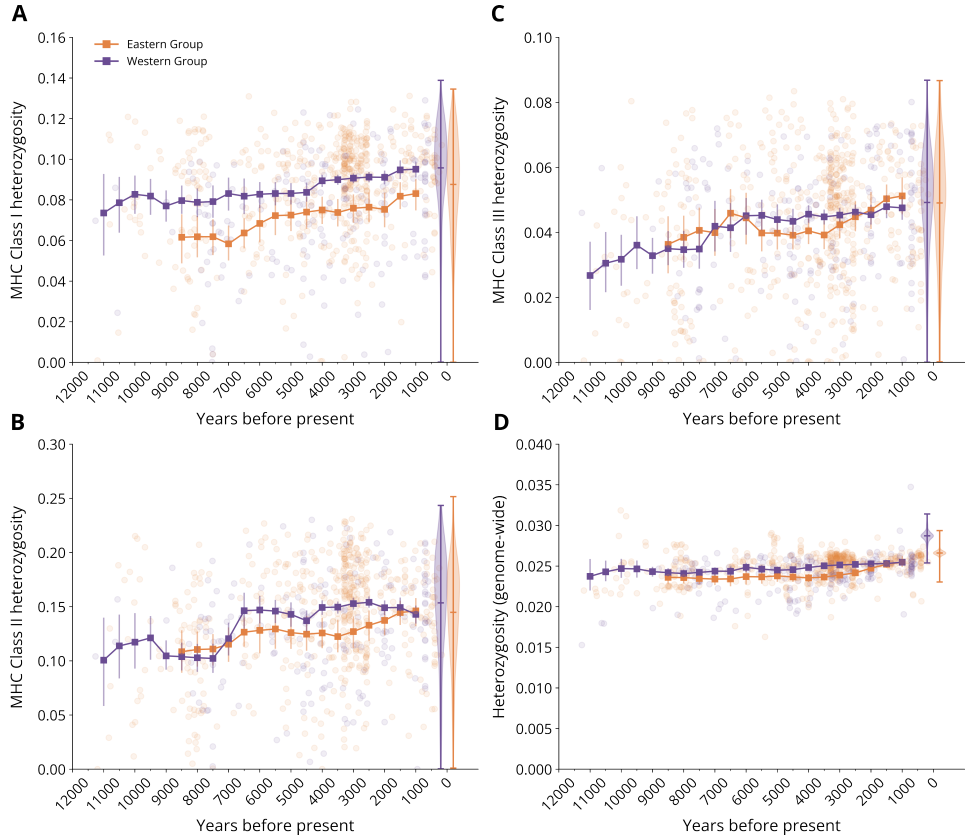


Figure S11: **Temporal trends of heterozygosity in the context of modern individuals from the Human Genome Diversity Project and 1000 Genomes Project**. Genomes were selected from the gnomAD panel that are geographically associated with the Western and Eastern groups (for a list of genomes used in this analyses, see `SI_table1.xlsx`). 2000-year sliding window of heterozygosity for the whole genome (A), MHC class I (B), MHC class II (C) and MHC class III (D), in the Western group (purple) and Eastern group (orange). Error bars indicate the 95% confidence intervals of the mean. Windows with less than 10 genomes are not shown.

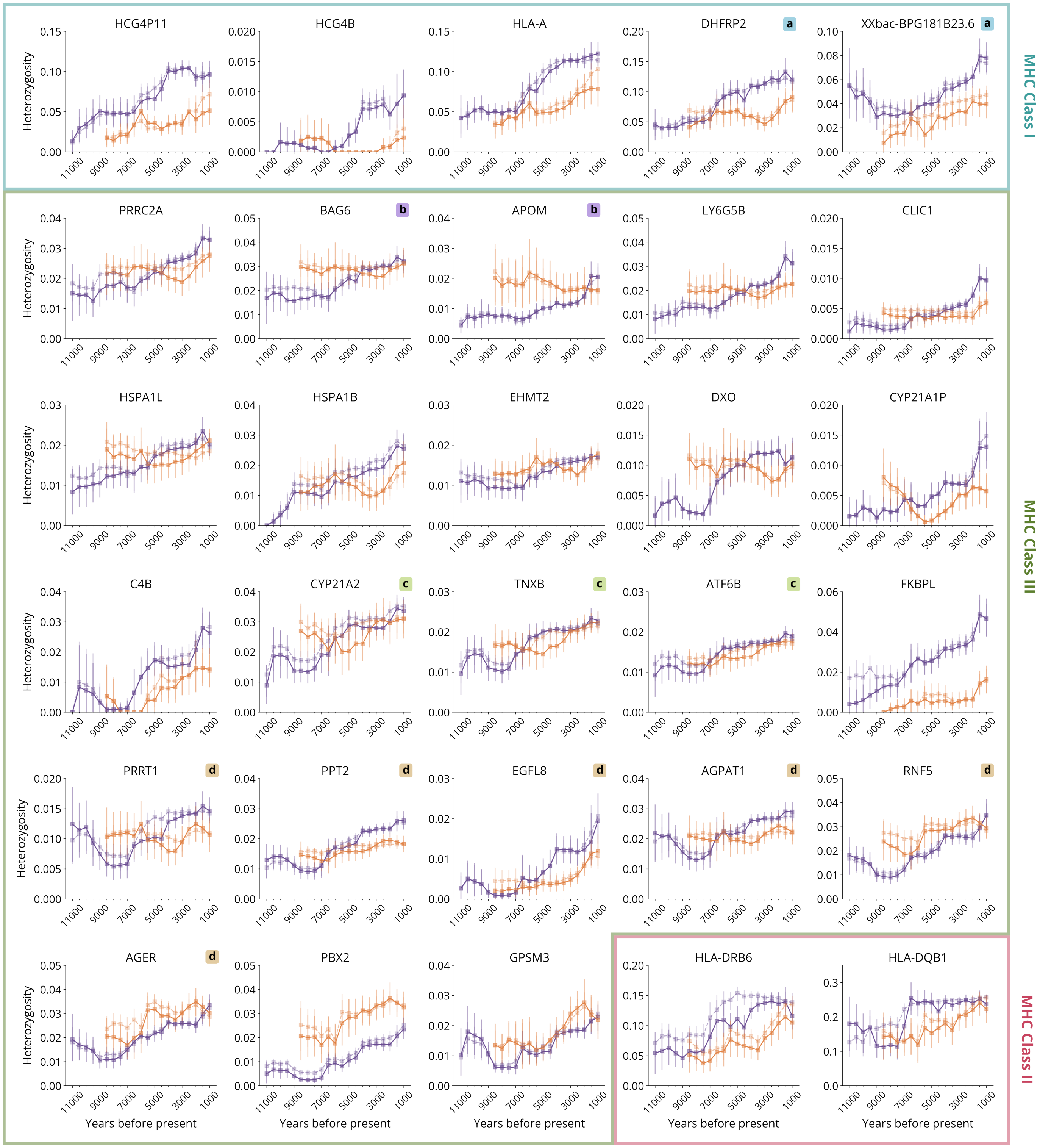


Figure S12: **Temporal trends of observed and expected heterozygosity for significant MHC genes from the Western group genomic scan**. Each panel shows the temporal trajectory of observed heterozygosity (solid lines) and expected heterozygosity (dashed lines) for the Western group (purple lines) and Eastern group (orange lines). Error bars show the 95% confidence intervals, generated by bootstrapping. Groups of genes that either overlap or exhibit linkage disequilibrium (LD) above a specified threshold in a modern LD panel ($\bar{r^2} \geq 0.2$) are labeled using lowercase letters placed in the top-right corner of each panel.

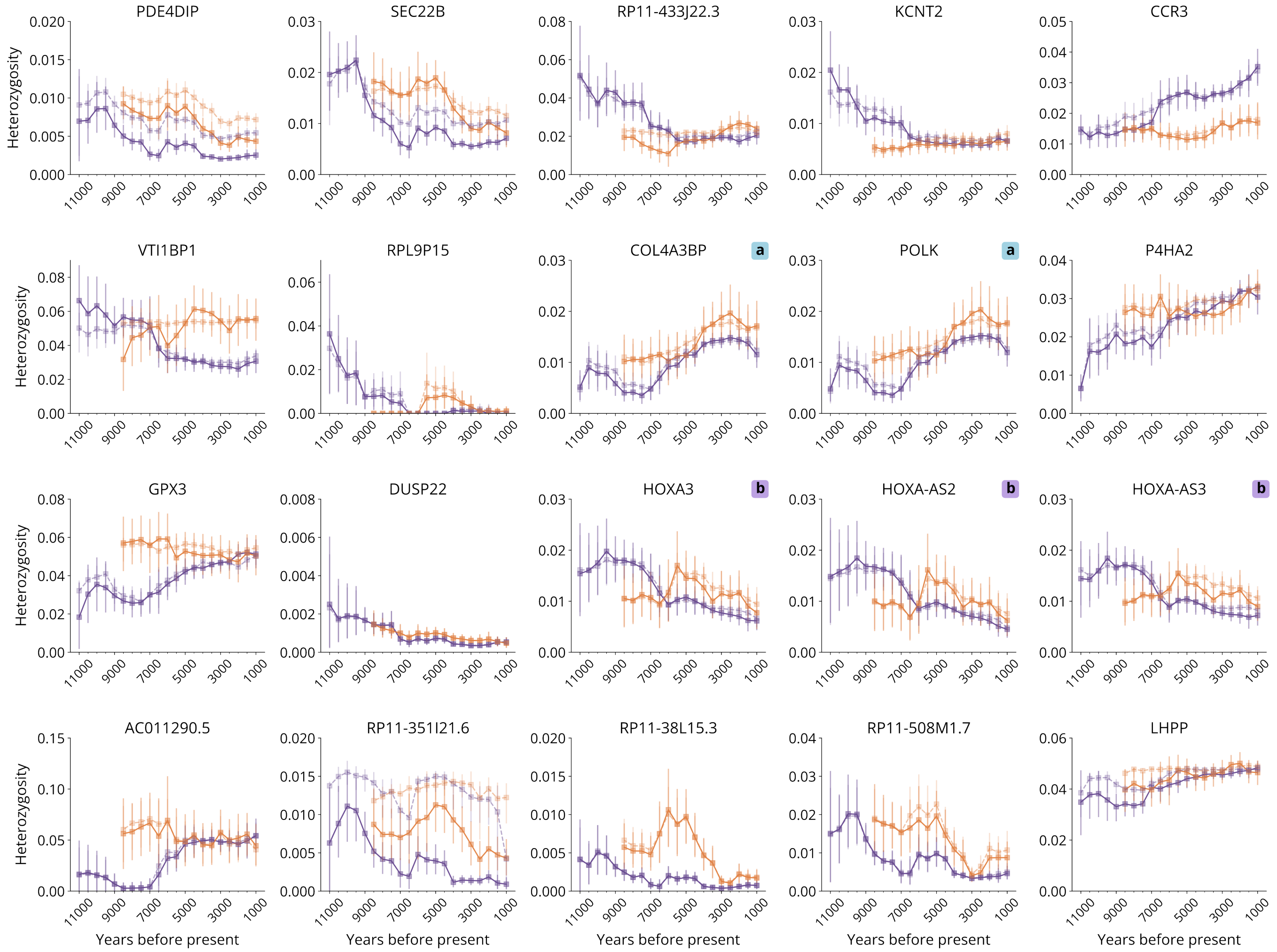


Figure S13: **Temporal trends of observed and expected heterozygosity for significant non-MHC genes from the Western group genomic scan**. Each panel shows the temporal trajectory of observed heterozygosity (solid lines) and expected heterozygosity (dashed lines) for the Western group (purple lines) and Eastern group (orange lines). Error bars show the 95% confidence intervals, generated by bootstrapping. Groups of genes that either overlap or exhibit linkage disequilibrium (LD) above a specified threshold in a modern LD panel ($\bar{r^2} \geq 0.2$) are labeled using lowercase letters placed in the top-right corner of each panel.

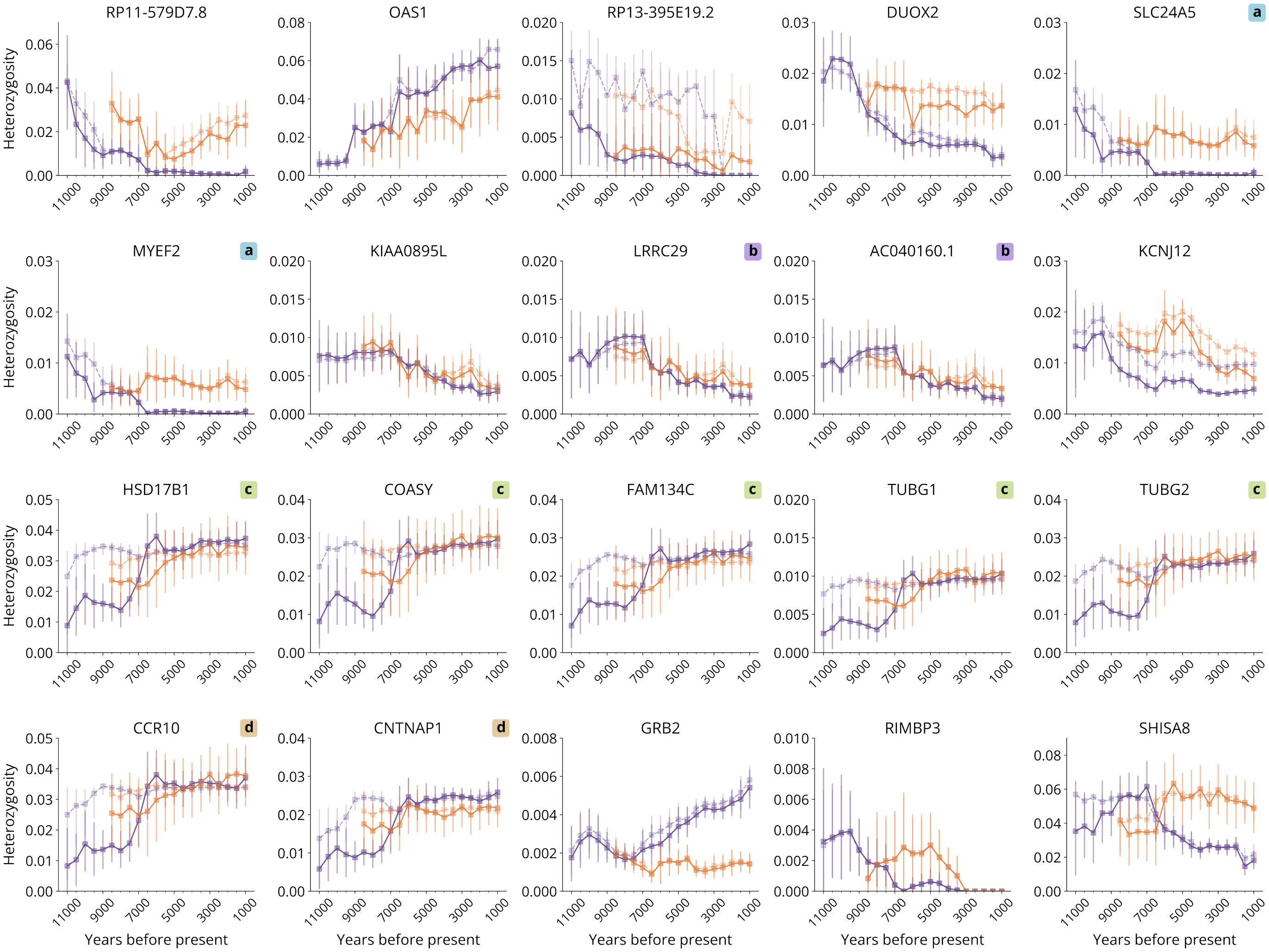


Figure S14: **Temporal trends of observed and expected heterozygosity for significant non-MHC genes from the Western group genomic scan**. Each panel shows the temporal trajectory of observed heterozygosity (solid lines) and expected heterozygosity (dashed lines) for the Western group (purple lines) and Eastern group (orange lines). Error bars show the 95% confidence intervals, generated by bootstrapping. Groups of genes that either overlap or exhibit linkage disequilibrium (LD) above a specified threshold in a model LD panel ($\bar{r^2} \geq 0.2$) are labeled using lowercase letters placed in the top-right corner of each panel.

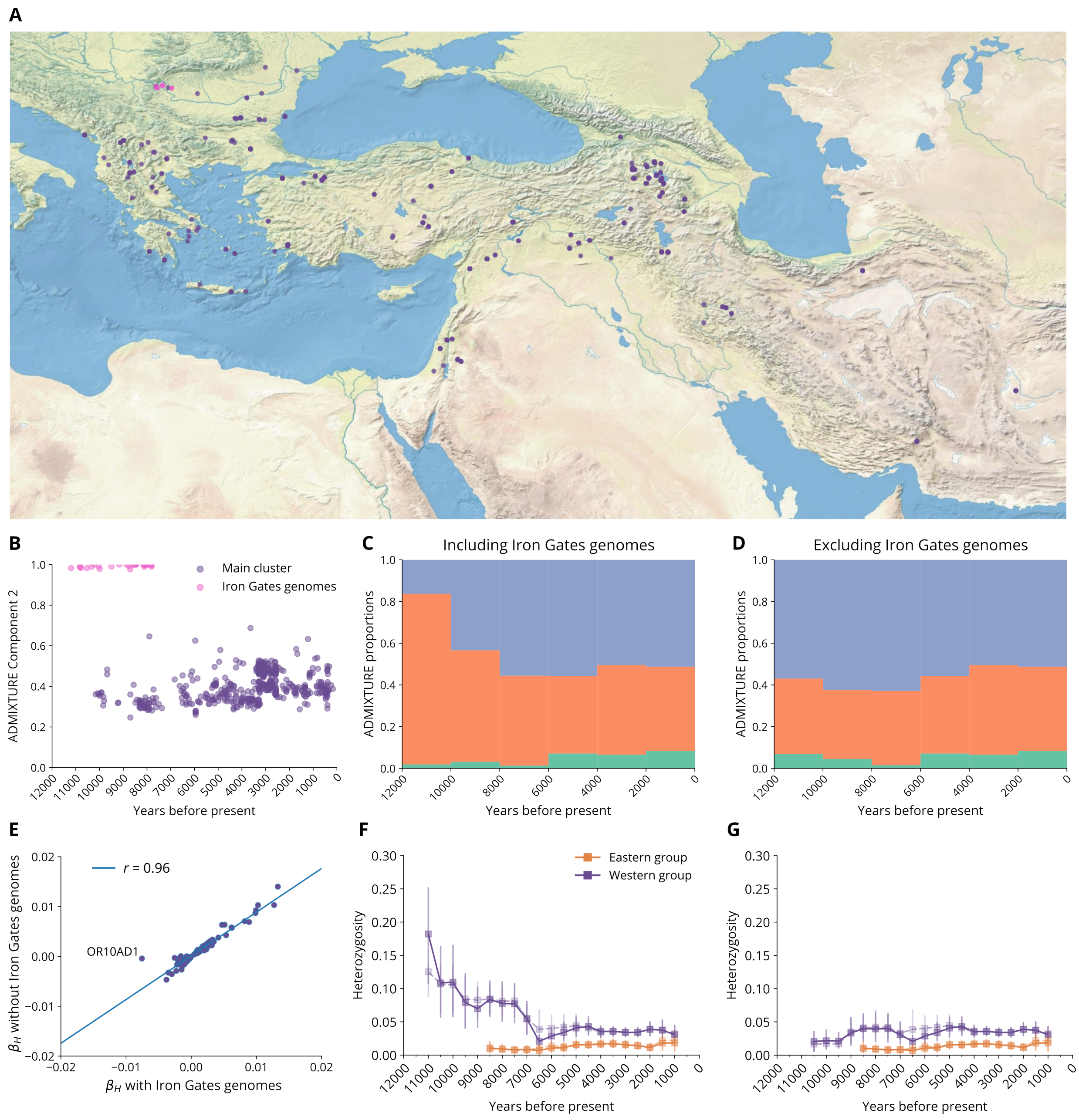


Figure S15: **Robustness of genomic scan to exclusion of early individuals from the Iron Gates region in the Western group**. In the Western group, the period between 12000–7000BP includes 29 individuals from the Iron Gates region. As these genomes cluster separately in our ancestry analysis and comprise 34% of the individuals of the Western group in this period, we assessed the robustness of our genomic scans to their exclusion. (A) Geographic positions of individuals included in the Western group, with the main cluster show as purple circles, and individuals in the Iron Gates region show as pink circles. (B) Distribution of the second ancestry component for genomes in the Western group taken from an ADMIXTURE model including all genomes in the AADR ($K = 3$) [33]. The Iron Gates genomes (pink circles) are distinct from other genomes in the group (purple circles). (C–D) ADMIXTURE result ($K = 3$) for genomes in the Western group including the Iron Gates genomes (C) and excluding the Iron Gates genomes (D), binned in 2000-year windows, showing the change in ancestry components over time. (E) Correlation between the results of the genomic scan for all genes in the Western group, including and excluding the Iron Gates individuals (pink circles in B). The outlier gene OR10AD1 is annotated. (F–G) Temporal trend of the heterozygosity of the outlier gene OR10AD1 in the Western group (purple) and Eastern group (orange), including the Iron Gates genomes (F) and excluding the Iron Gates genomes (G). Dashed lines indicate the expected heterozygosity, computed from the allele frequencies for all samples in each window. Vertical bars indicate 95% CIs of the mean.
